# Shuffled ATG8 interacting motifs form an ancestral bridge between UFMylation and C53-mediated autophagy

**DOI:** 10.1101/2022.04.26.489478

**Authors:** Lorenzo Picchianti, Víctor Sánchez de Medina Hernández, Ni Zhan, Nicholas A. T. Irwin, Madlen Stephani, Harald Hornegger, Rebecca Beveridge, Justyna Sawa-Makarska, Thomas Lendl, Nenad Grujic, Sascha Martens, Thomas A. Richards, Tim Clausen, Silvia Ramundo, G. Elif Karagöz, Yasin Dagdas

## Abstract

UFMylation mediates the covalent modification of substrate proteins with UFM1 (Ubiquitin-fold modifier 1) and regulates the selective degradation of endoplasmic reticulum (ER) via autophagy (ER-phagy) to maintain ER homeostasis. Specifically, collisions of the ER-bound ribosomes trigger ribosome UFMylation, which in turn activates C53-mediated autophagy that clears the toxic incomplete polypeptides. C53 has evolved non-canonical shuffled ATG8 interacting motifs (sAIMs) that are essential for ATG8 interaction and autophagy initiation. Why these non-canonical motifs were selected during evolution, instead of canonical ATG8 interacting motifs remains unknown. Here, using a phylogenomics approach, we show that UFMylation is conserved across the eukaryotes and secondarily lost in fungi and some other species. Further biochemical assays have confirmed those results and showed that the unicellular algae, *Chlamydomonas reinhardtii* has a functional UFMylation machinery, overturning the assumption that this process is linked to multicellularity. Our conservation analysis also revealed that UFM1 co-evolves with the sAIMs in C53, reflecting a functional link between UFM1 and the sAIMs. Using biochemical and structural approaches, we confirmed the interaction of UFM1 with the C53 sAIMs and found that UFM1 and ATG8 bound to the sAIMs in a different mode. Conversion of sAIMs into canonical AIMs prevented binding of UFM1 to C53, while strengthening ATG8 interaction. This led to the autoactivation of the C53 pathway and sensitized *Arabidopsis thaliana* to ER stress. Altogether, our findings reveal an ancestral toggle switch embodied in the sAIMs that regulates C53-mediated autophagy to maintain ER homeostasis.

## Introduction

Perturbations of cellular homeostasis, termed “cellular stress”, triggers protein aggregation and impairment of organelle function, and reduces organismal fitness and lifespan. Quality control pathways closely monitor the health of cellular components to alleviate cellular stress [1]. Cells first try to rescue aberrant proteins and organelles to restore cellular homeostasis [2–4]. If these attempts fail, dysfunctional proteins and organelles are rapidly degraded [5]. Defects in cellular quality control has been linked to several diseases, including cognitive decline, aging, cancer, and metabolic disorders in humans, and reduced stress tolerance and fitness in plants [1, 6–8]. Although, studies in the last decade have revealed a comprehensive suite of interconnected pathways that mediate protein and organelle degradation, the regulatory mechanisms that keep them switched off under normal conditions remain largely unknown.

Selective autophagy is a major quality control pathway that degrades unwanted or harmful cellular components including protein aggregates or damaged organelles with high precision [9]. Modular selective autophagy receptors (SARs) bring those cargo to the core autophagy machinery, resulting in their selective degradation [8, 10]. SARs recruit the autophagy machinery through their interaction with ATG8, a ubiquitin-like protein conjugated to the phagophore, and ATG11/FIP200, a scaffold protein of the autophagy initiation complex ATG1/ULK1 [11]. Recent structure-function studies have shown that SARs interact with ATG8 via various amino acid sequence motifs [4]. The **c**anonical **A**TG8 **I**nteracting **M**otif, (**cAIM**), also known as an **L**C3 **I**nteracting **R**egion (**LIR**), is a well characterized short linear motif that interacts with ATG8 by forming a parallel β-sheet with the β-sheet 2 in ATG8 [12]. The cAIM is represented by the WXXL consensus sequence, where W is an aromatic residue (W/F/Y), L is a aliphatic hydrophobic residue (L/I/V), and X can be any residue [13]. Recently, we showed that the ER-phagy receptor C53 (CDK5RAP3 in humans) interacts with plant and mammalian ATG8 isoforms via a non-canonical AIM sequence, with the consensus sequence IDWG/D, which we named the shuffled AIM (sAIM) [14]. However, the structural basis of sAIM-ATG8 interaction and its importance in C53-mediated autophagy and endoplasmic reticulum homeostasis remain unknown.

Our work and a recent genome wide CRISPR screen revealed that selective ER autophagy (ER-phagy) is regulated by UFMylation [14, 15]. UFMylation is similar to ubiquitination, where UFM1 is conjugated to substrate proteins via an enzymatic cascade [16, 17]. First, UFM1 is cleaved to its mature form by the protease UFSP2. UFM1 is then activated by UBA5, an E1 activating enzyme. UBA5 transfers UFM1 to UFC1, the E2 conjugating enzyme, through a trans-binding mechanism [18, 19]. Finally, UFM1 is transferred to the substrate by UFL1, which, in complex with the ER membrane protein DDRGK1, form an E3 ligase complex to covalently modify lysine residues on substrates [20, 21]. To date, the best characterized UFMylation substrate is the 60S ribosomal subunit RPL26 [22]. RPL26 UFMylation is triggered by stalling of ER-bound ribosomes and is necessary for autophagic degradation of the incomplete polypeptides trapped on ER-bound ribosomes [15, 23]. We have shown that C53 mediates the degradation of these incomplete polypeptides in a UFMylation-dependent manner [14]. However, how UFMylation regulates C53-mediated autophagy remains unknown.

Here, we combined evolutionary analyses with cellular and structural biology experiments to investigate the regulation of C53-mediated autophagy via UFMylation. We reconstructed the evolutionary history of the UFMylation pathway and found that it is ubiquitous across eukaryotes, suggesting its presence in the last eukaryotic common ancestor. Based on our phylogenetic analyses, we reconstituted the UFMylation machinery of the unicellular green algae, *Chlamydomonas reinhardtii*, and showed that it is functional and essential for the ER-stress tolerance, demonstrating the importance of UFMylation beyond plants and animals. Biochemical and structural studies, supported with evolutionary correlation analyses revealed that shuffled AIMs (sAIMs) within C53 intrinsically disordered region (IDR) form versatile binding sites that allow C53 to interact with both ubiquitin-like proteins (UBLs), UFM1 and ATG8. However, ATG8 and UFM1 bind these motifs in a different mode. While ATG8 bound strongest to cAIM and displayed equal preference for the first and the second sAIM in C53 IDR, UFM1 interacted preferentially with the first sAIM. Conversion of sAIMs in C53 into canonical AIMs shifted its binding preference towards ATG8 and led to premature activation of autophagy driven by C53, sensitizing *Arabidopsis thaliana* to ER stress. Altogether, our findings reveal an ancient UFM1 dependent regulatory mechanism that prevents premature activation of C53-mediated autophagy.

## Results

### The UFMylation pathway is conserved across eukaryotes and functional in the unicellular alga, *Chlamydomonas reinhardtii*

To explore a potential link between the UFMylation pathway and C53-mediated autophagy, we searched for the existence of proteins involved in UFMylation across the eukaryotic tree of life using a phylogenomic approach across 151 species. We identified the presence of UFMylation proteins in all major eukaryotic lineages, indicating that the UFMylation pathway was a feature of the last eukaryotic common ancestor (Fig. 1A, Fig. S1). Despite its ancestral origin, multiple groups have lost parts or all the UFMylation proteins. Apparent absence of a gene family can result from dataset incompleteness (e.g., incomplete genome assembly and annotation) but recurrent absences across multiple closely related genomes is strong evidence that a protein has been lost from those genomes and specific branches of the tree of life. We noted the loss of UFMylation from multiple parasitic and algal lineages as well as in fungi (Fig. 1A). Gene loss in parasites is a recurrent phenomenon, resulting from parasitic genome streamlining [24], but the absence of UFMylation in genera such as *Plasmodium*, *Entamoeba*, and *Trichomonas* indicates that the pathway is often expendable in parasitic organisms (Fig. 1A). UFMylation has also been lost repeatedly in algal lineages, suggesting that life history or other shared cellular characters may dictate the pathway’s retention. Similar to parasites and algae, fungi have also lost UFMylation, although certain lineages retain pathway components, indicating that either repeated losses have occurred, or genes were lost and subsequently reacquired through horizontal gene transfer (Fig. 1A). Lastly, despite the loss of UFM1 in various lineages, certain UFMylation pathway proteins are occasionally retained, particularly DDRGK1, UFL1, and in a few cases, C53 (e.g., the oomycete genus *Albugo* and the chytrid class Neocallimastigomycetes) (Fig. 1A). This suggests that these proteins may have additional cellular functions independent of UFM1. Altogether, these data demonstrate that the UFMylation pathway is present throughout eukaryotes, implying that it is functionally conserved in both unicellular and multicellular species, unlike suggested before [22].

**Figure 1.**
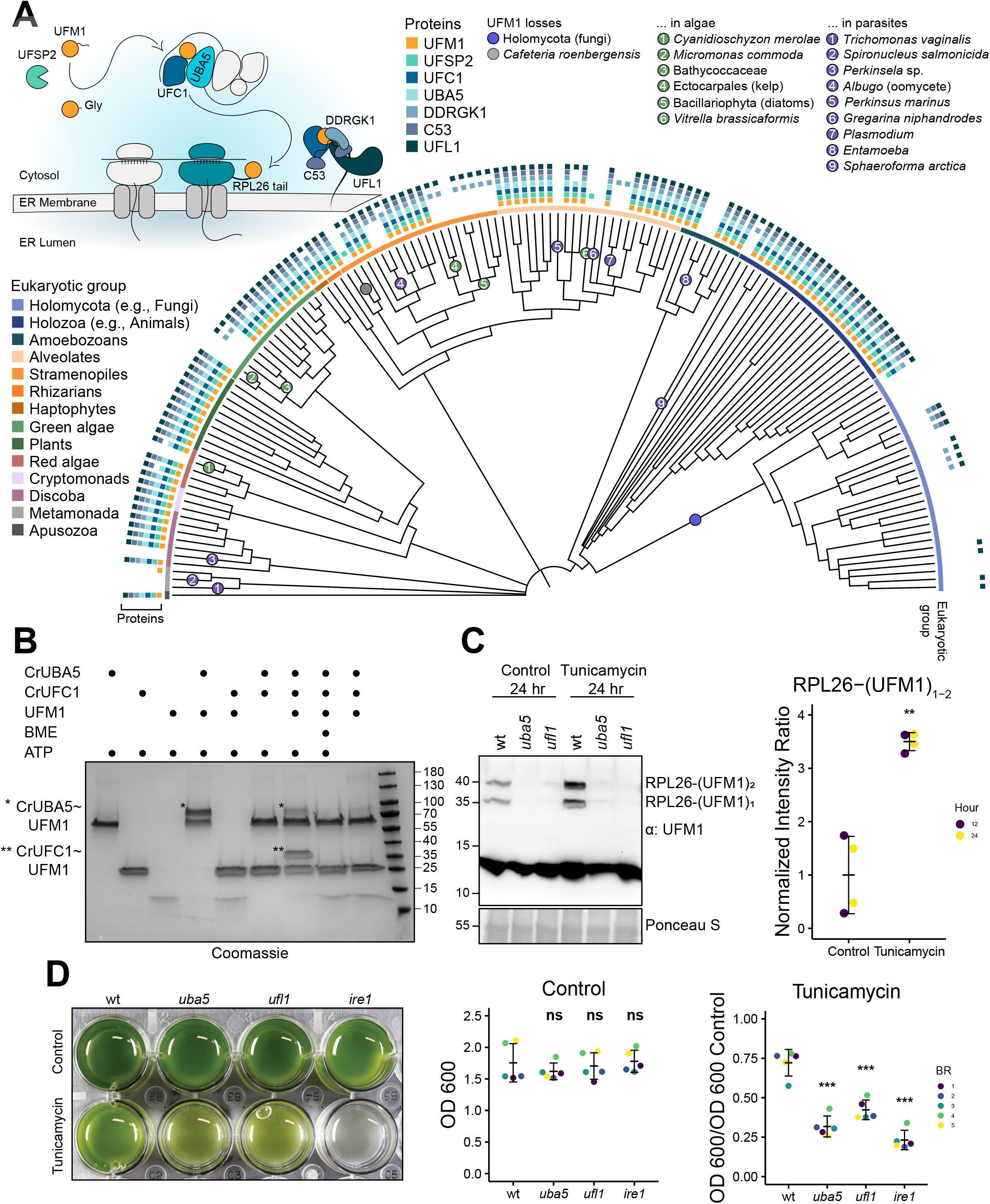
UFMylation did not evolve in multicellular eukaryotes. **(A) A eukaryotic phylogeny displaying the presence or absence of UFMylation proteins across diverse species.** Protein presence is displayed at the tip of each tree and major eukaryotic taxonomic groups are denoted with a colored ribbon. Losses of UFM1 have been highlighted. A schematic diagram depicting UFMylation cascade and C53-receptor complex has been included for reference. See Fig. S1 for an expanded phylogeny, including species names. **(B) *Chlamydomonas reinhardtii* (Cr) UBA5 and UFC1 are active E1 and E2 enzymes.** SDS-PAGE analysis showing transfer of UFM1 to CrUBA5 and CrUFC1. The gels are run in non-reducing conditions except where otherwise specified. The presented gel is representative of two independent experiments. BME: β-mercaptoethanol; ATP: Adenosine triphosphate. **(C) RPL26 mono- and di-UFMylation is lost in *Chlamydomonas reinhardtii* (Cr) *uba5* and *ufl1* mutants.** Liquid TAP cultures were either left untreated (control) or treated for 24 hours with 200 ng/mL tunicamycin. Protein extracts were analyzed by immunoblotting with anti-UFM1 antibodies. Total proteins were analyzed by Ponceau S staining. 12 hours and 24 hours treatment replicates are shown in Fig. S3C. *Right Panel*, Quantification of UFMylated RPL26. Bars represent the mean (± SD) of 2 biological replicates. Two-tailed unpaired t-test with Welch correction was performed to analyze the differences between control and treated samples. **, p-value < 0.01. RPL26-(UFM1)_1_: RPL26 mono-UFMylated; RPL26-(UFM1)_2_: RPL26 di-UFMylated. **(D) *Chlamydomonas reinhardtii* (Cr) UFMylation pathway mutants are sensitive to ER stress triggered by tunicamycin.** Liquid TAP cultures of wild type (wt), *uba5, ufl1* and *ire1* mutants were either left untreated (control) or treated for 3 days with 200 ng/mL of tunicamycin. *Left panel,* representative images of control and treated liquid cultures taken 3-days after incubation. *Middle Panel,* optical density (OD) 600 (OD_600_) quantification of each genetic background under control conditions. Bars represent the mean (± SD) of 5 biological replicates. Two-tailed unpaired t-tests were performed to analyze the differences between wild type and mutants. *Right Panel,* normalized OD_600_ quantification of each genetic background under tunicamycin treatment conditions. Bars represent the mean (± SD) of 5 biological replicates. Two-tailed unpaired t-tests were performed to analyze the differences between wild type and mutants. ns, p-value > 0.05; ***, p-value < 0.001. BR: Biological Replicate.

To characterize the functionality of the UFMylation pathway in a unicellular species, we investigated UFMylation in *Chlamydomonas reinhardtii* (Cr), a single-celled green alga. We purified CrUBA5, CrUFC1 and CrUFM1 and tested their ability to conjugate UFM1. *In vitro* E2-charging of CrUFM1 worked similar to the human UFMylation cascade [18]. In a UBA5-dependent manner, UFM1 was transferred to UFC1 by formation of a thioester bond, which could be reduced by β-mercaptoethanol (Fig. 1B). This indicates that the UFM1 conjugation mechanism is conserved in *C. reinhardtii*, prompting us to tested substrate UFMylation. We first examined conservation of the RPL26 tail, which has been shown to be ufmylated [22]. Protein sequence alignment and Twincons analysis revealed that the ufmylated lysine residues in RPL26 are conserved in species with UFM1, including *C. reinhardtii* (Fig. S2). Moreover, immunoblot analysis using a UFM1 antibody revealed two bands corresponding to mono- and di-ufmylated RPL26 (Fig. 1C). RPL26 UFMylation was dependent on the UFMylation machinery, as both bands were absent in *uba5* and *ufl1* mutants (Fig. 1C, Fig. S3). Consistent with previous studies [14, 23], RPL26 UFMylation was induced upon ER stress triggered by tunicamycin, a glycosylation inhibitor that leads to the accumulation of unfolded proteins in the ER (Fig. 1C). Finally, we performed ER stress tolerance assays to test the physiological importance of UFMylation in *C. reinhardtii*. *uba5* and *ufl1* mutants were more sensitive to ER stress than the wild type, confirming UFMylation is essential for ER stress tolerance in *C. reinhardtii* (Fig. 1D). Altogether, these findings suggest UFMylation contributes to ER homeostasis across eukaryotes.

### C53 interacts with UFM1 via the shuffled ATG8 interacting motifs (sAIMs)

In addition to revealing the conservation of the UFMylation pathway in unicellular organisms, our phylogenomic analysis also showed a strong presence-absence correlation between C53 and UFM1 (Fig. 1A). To investigate whether this correlation is due to a functional link between C53 and UFM1, we first performed ConSurf analysis of C53 to estimate the conservation of each residue [25]. C53 has two α-helical domains at the N-and C-termini, connected with an intrinsically disordered region. In contrast to the alpha helical domains, which were highly conserved, the IDR was divergent. However, within the IDR, there were four highly conserved regions that corresponded to the sAIMs (Fig. 2A). To explore a possible connection between UFM1 and the sAIMs, we examined the conservation of individual sAIMs between species with and without UFM1 (Fig. 2B). Although IDR residues are generally not conserved between and within groups, the sAIMs show a strong dichotomy between species with and without UFM1, demonstrating a link between sAIM conservation and the presence of UFM1. In agreement with this, multiple sequence alignment revealed that the C53 IDRs in species lacking UFM1 are consistently shorter relative to UFM1-encoding species, and lack sAIMs (Fig. 2C). To support these findings, we synthesized C53 homologs from two species that lack UFM1 (the oomycete *Albugo candida* (Ac) and chytrid *Piromyces finnis* (Pf)) and tested whether they interact with UFM1 or ATG8 using *in vitro* pulldown assays. Both AcC53 and PfC53 were able to interact with Arabidopsis ATG8A and human ATG8 isoform GABARAP (Fig. 2D), but they did not interact with either of the UFM1 orthologs tested (Fig. 2E). Their ability to bind ATG8 may be due to the presence of putative cAIMs within the truncated IDRs of both AcC53 and PfC53 (Fig. 2C, D).

**Figure 2.**
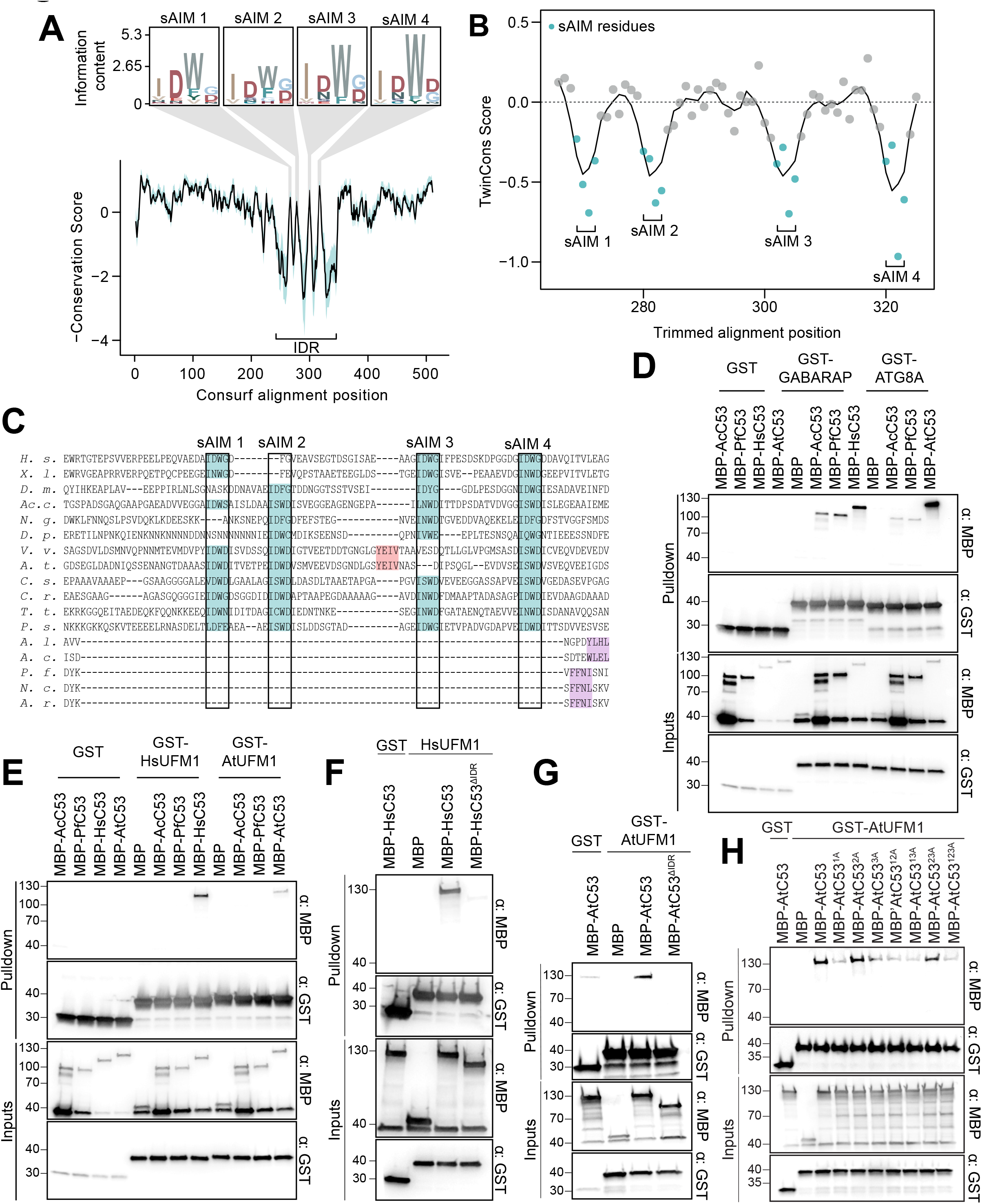
The sAIM sequences within C53 Intrinsically Disordered Region (IDR) are highly conserved and essential for UFM1 interaction. **(A) ConSurf conservation analysis of C53 from diverse eukaryotes.** Conserved regions within the IDR (intrinsically disordered region) have been highlighted and supplemented with sequence logos. **(B) TwinCons analysis comparing the conservation and divergence of C53 among species with and without UFM1.** The four regions corresponding to the sAIMs have been highlighted. Negative values reflect divergent signature regions between the two species groups. (**C) A trimmed multiple sequence alignment depicting the conservation of the sAIMs.** The four sAIMs and cAIMs in plants and UFM1-lacking species have been highlighted in teal and light red, respectively. Putative cAIMs are highlighted in purple. Abbreviations: *H. s.*, *Homo sapiens*; *X. l., Xenopus laevis; D. m., Drosophila melanogaster; Ac. c., Acanthamoeba castellanii; N. g., Naegleria gruberi*; *D. p., Dictyostelium purpurea; V. v., Vitis vinifera; A. t., Arabidopsis thaliana* (trimmed sequence)*; C. s., Chlorella sorokiniana; C. r., Chlamydomonas reinhardtii; T. t., Tetrahymena thermophila; P. s. Phytopthora sojae; A. l., Albugo laibachii; A. c., Albugo candida; P. f., Piromyces finnis, N. c., Neocallimastix californiae; A. r., Anaeromyces robustus*. **(D, E) AcC53 and PfC53 do not have sAIM sequences and cannot interact with UFM1**. Ac: *Albugo candida*, Pf: *Piromyces finnis*. **(F, G) C53 IDR is essential for UFM1 interaction**. HsC53 (B) and AtC53 (C) IDRs are necessary to mediate the interaction with AtUFM1 and HsUFM1 respectively. MBP-AtC53^ΔIDR^: MBP-AtC53^(1-239, (KGSGSTSGSG)2, 373-549)^; MBP-HsC53^ΔIDR^: HsC53^(1-262, (KGSGSTSGSG), 317-506)^. **(H) AtC53^sAIM^ cannot interact with AtUFM1.** Individual or combinatorial mutations in sAIM1 (1A: W276A), sAIM2 (2A: W287A) and sAIM3 (3A: W335A) suggest sAIM1 is crucial for UFM1 interaction. (B, C, E, F, G) Bacterial lysates containing recombinant protein were mixed and pulled down with glutathione magnetic agarose beads. Input and bound proteins were visualized by immunoblotting with anti-GST and anti-MBP antibodies.

As the phylogenomic analyses suggested that the sAIMs have been retained to mediate C53-UFM1 interaction, we sought to reconstitute the human UFM1-C53 complex using native Mass-Spectrometry (nMS). We found that C53 binds to human UFM1 in a 1:1 or 1:2 stoichiometry, similar to the C53-GABARAP interaction (Fig. S4). To map the UFM1 interacting region in C53, we performed *in vitro* pulldowns with *Homo sapiens* (Hs) and *Arabidopsis thaliana* (At) C53 truncations. As in the C53-ATG8 interaction, the C53 IDR was necessary for interaction between C53 and UFM1 (Fig. 2F, G). Further individual and combinatorial mutagenesis of the tryptophan residues in sAIMs showed that the UFM1-C53 interaction is mediated by sAIMs located in the IDR (Fig. 2H).

We next asked whether ATG8 and UFM1 bind the sAIMs in a similar manner. First, we performed nMS analysis to test the interaction of HsUFM1 with a canonical AIM (cAIM) peptide [14]. Unlike the UBA5-LIR peptide (GPLHDDNEWNISVVDD), which has been shown to interact with UFM1 [26–28], the cAIM peptide did not appreciably interact with UFM1 (Fig. 3A). Consistently, the cAIM peptide outcompeted the GABARAP-C53 interaction but not the HsUFM1-C53 interaction (Fig. 3B). *C. reinhardtii* proteins behaved similarly; CrC53 interacted with ATG8 in a cAIM-dependent manner and CrUFM1 in a cAIM-independent manner (Fig. S5).

**Figure 3.**
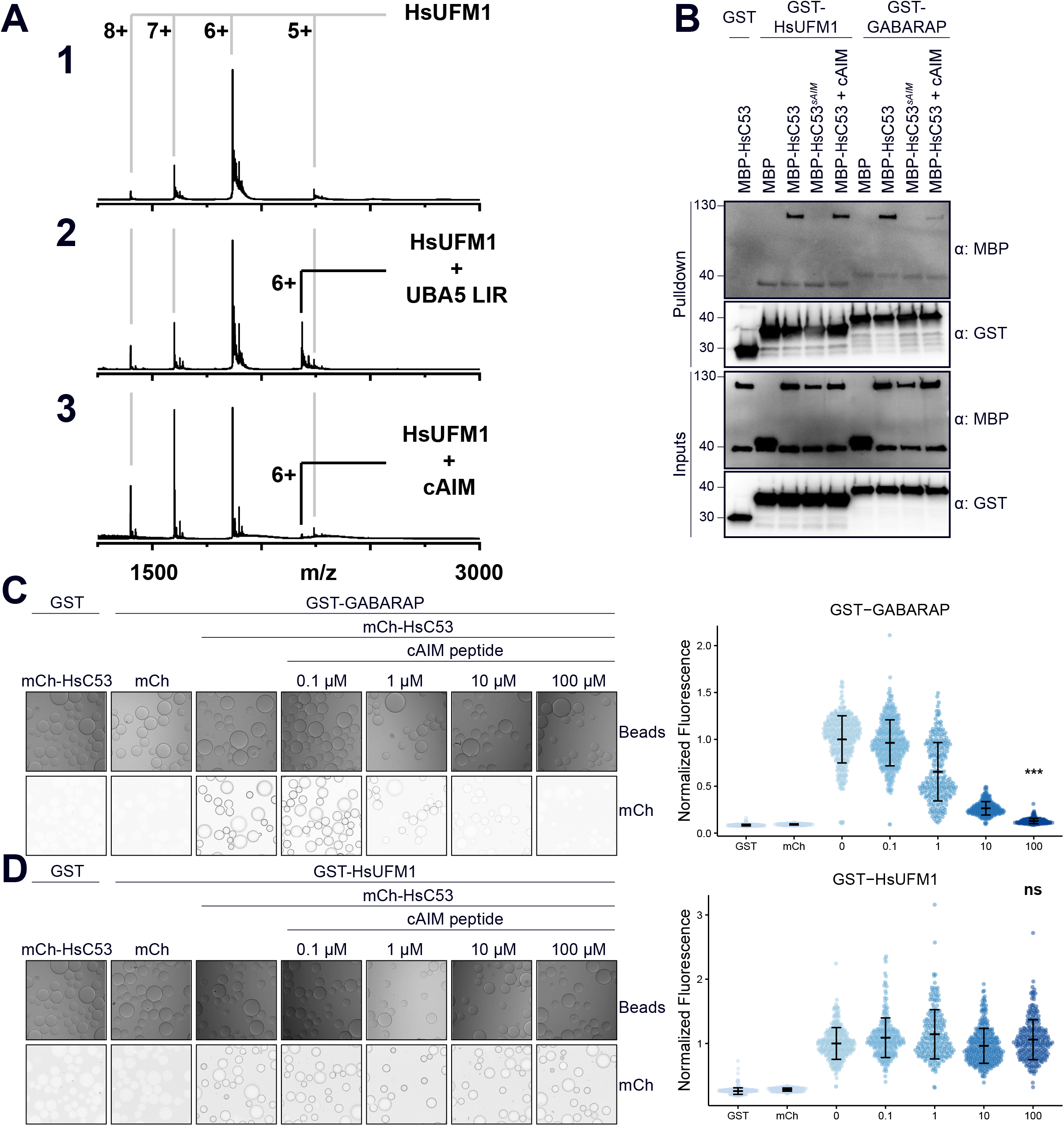
The canonical ATG8 Interacting Motif (cAIM) cannot outcompete C53-UFM1 interaction. **(A) Complex formation between cAIM peptide and UFM1.** Native mass spectrometry (nMS) spectra of (1) HsUFM1 (5 µM), (2) HsUFM1 (5 µM) and UBA5 LIR peptide (25 µM) and (3) HsUFM1 (5 µM) and cAIM peptide (25 µM). UFM1 forms a 1:1 complex with the UBA5 LIR peptide. Only a negligible amount of 1:1 complex is formed between the cAIM peptide and UFM1, indicating a lower affinity interaction. **(B) The cAIM peptide cannot outcompete HsUFM1-HsC53 interaction.** Bacterial lysates containing recombinant protein were mixed and pulled down with glutathione magnetic agarose beads. Input and bound proteins were visualized by immunoblotting with anti-GST and anti-MBP antibodies. cAIM peptide was used to a final concentration of 200 µM. HsC53^sAIM^: HsC53^W269A, W294A, W312A^. **(C, D) Microscopy-based protein–protein interaction assays showing unlike GABARAP-C53 interaction, UFM1-C53 interaction is insensitive to cAIM peptide competition**. Glutathione-sepharose beads were prepared by incubating them with GST-GABARAP (C) or GST-HsUFM1 (D). The pre-assembled beads were then washed and mixed with 1 µM of HsC53 containing increasing concentrations of cAIM peptide (0-100 µM). The beads were then imaged using a confocal microscope. *Left Panel,* representative confocal images (inverted grayscale) for each condition are shown. *Right panel*, normalized fluorescence is shown for each condition with the mean (± SD) of 4 replicates. Unpaired two-samples Wilcoxon test with continuity correction was performed to analyze the differences between wild type and wild type with 100 µM AIM peptide. ns, not significant, p-value > 0.05, ***, p-value < 0.001. Total number of beads, mean, median, standard deviation and p-values are reported in Supplementary data 2.

To further test these interactions, we performed microscopy-based on-bead binding assays. The advantage of this technique is the ability to visualize protein-protein interactions with fast dissociation constants at equilibrium. It can also detect relatively weak, transient interactions [29]. We purified GST-tagged Arabidopsis and human ATG8 and UFM1 proteins and coupled them to the glutathione coated beads (Sepharose 4B, Cytiva). We then tested whether mCherry tagged Arabidopsis and human C53 proteins could bind to the ATG8 or UFM1 coupled beads (Fig. S6A). Arabidopsis and human C53 interacted with wild type ATG8 and UFM1, and HsC53-GABARAP and AtC53-ATG8A interaction was outcompeted with increased concentrations of the cAIM peptide (Fig. 3C and Fig. S6B). In contrast, the cAIM peptide could not outcompete the HsC53-HsUFM1 or AtC53-AtUFM1 interaction (Fig. 3D and Fig. S6C). Consistently, the UBA5-LIR peptide and GABARAP were able to disrupt C53-UFM1 interaction (Fig. S6D). Altogether, these results suggested that ATG8 and UFM1 bind the sAIMs within C53 IDR, albeit in a different manner.

### Comparative NMR spectroscopy analysis revealed the differences between C53 IDR-UFM1 and C53 IDR-ATG8 interaction

To elucidate the difference between UFM1 and ATG8 binding to C53 IDR, we performed comparative nuclear magnetic resonance (NMR) spectroscopy analysis. We first obtained backbone resonance assignments of AtC53 IDR. We could assign 89% of the residues in AtC53 IDR. The sAIMs in AtC53 IDR share high sequence homology, therefore we validated the assignments using sAIM1 (AtC53 IDR^W276A^) and sAIM2 (AtC53 IDR^W287A^) mutants (Fig. 4A, Fig. S7A). The 2D heteronuclear single quantum correlation (HSQC) spectrum of ^15^N-labelled AtC53 IDR displayed small dispersion of the backbone amide residues, validating its intrinsically disordered nature. The NMR signals are sensitive to their chemical environment; binding of an interaction partner or conformational changes induced by protein-protein interaction shifts the NMR spectra. Moreover, NMR signal intensity drops mainly due to an increase in molecular weight upon complex formation and the chemical exchange that happens at the interaction surface [30–33].

**Figure 4.**
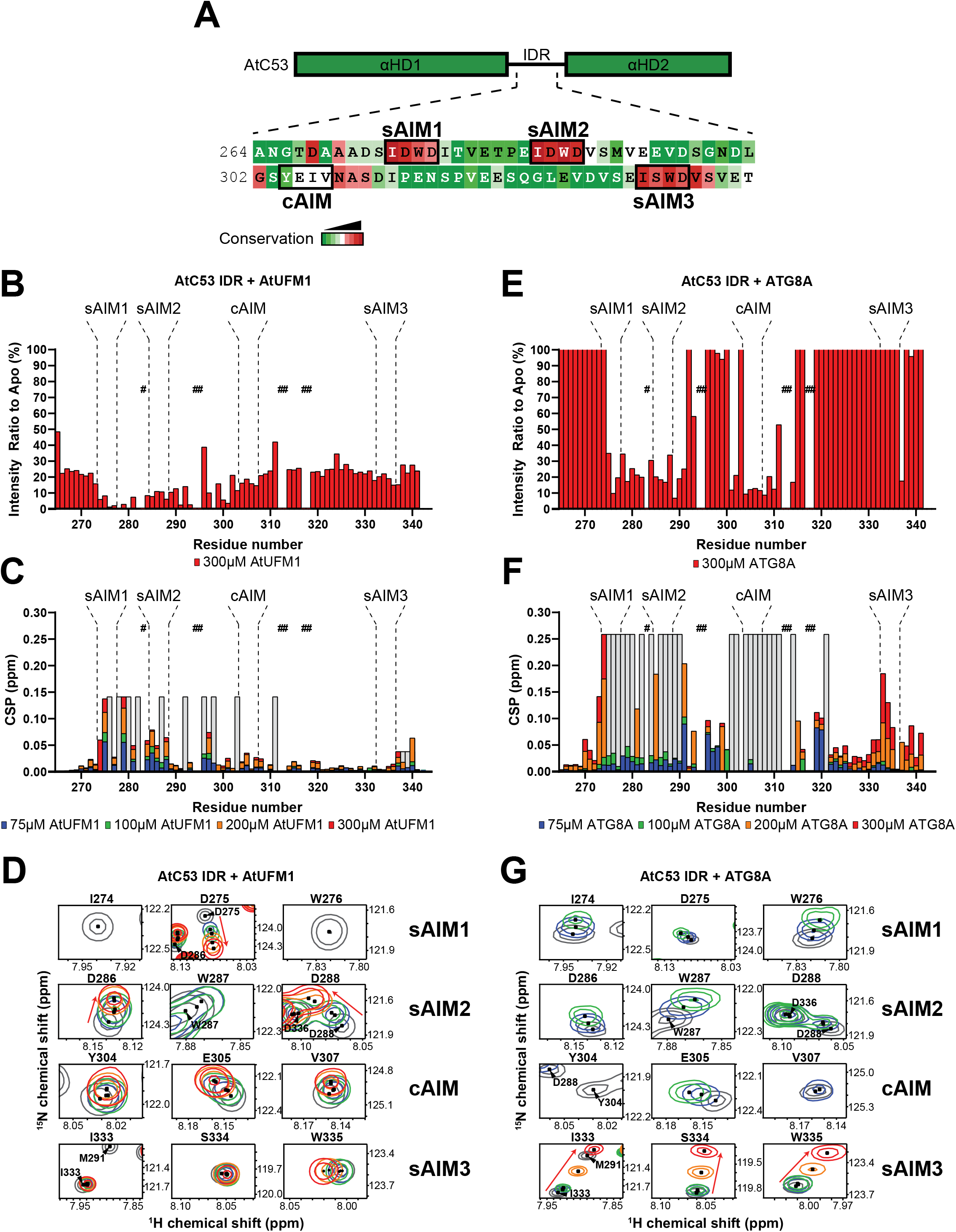
Comparative Nuclear Magnetic Resonance (NMR) spectroscopy analyses show C53 IDR-UFM1 interaction is different than C53 IDR-ATG8 interaction. **(A) AtC53 IDR harbours highly conserved canonical and shuffled ATG8 interaction motifs.** Schematic representation of AtC53 domains with the primary sequence of C53 IDR. The AIM sequences and their conservation are indicated with rectangular boxes and a color code, respectively. **(B) Binding of AtUFM1 to AtC53 IDR leads to a general drop in signal intensity.** Intensity ratio broadening of AtC53 IDR (100 µM) in the presence of 300 µM AtUFM1. Bars corresponding to residues in the AIMs are highlighted. **(C) UFM1-IDR binding involves sAIM1 and sAIM2.** NMR chemical shift perturbations (CSP) of AtC53 IDR (100 µM) in the presence of 75 µM (blue), 100 µM (green), 200 µM (orange) and 300 µM (red) AtUFM1. **(D) AtC53 IDR spectra signals shift upon AtUFM1 addition in a concentration-dependent manner.** Insets of overlaid ^1^H-^15^N HSQC spectra of isotope-labeled AtC53 IDR (100 µM) showing chemical shift perturbations of individual peaks from backbone amides of AIM residues in their free (gray) or bound state to unlabeled AtUFM1. Chemical shifts are indicated with arrows. **(E) Binding of ATG8A AtC53 IDR leads to a localized signal intensity drop in sAIM1-2 and cAIM regions.** Intensity ratio broadening of C53 IDR (100 µM) in the presence of 300 µM ATG8A. Bars corresponding to residues in AIMs are highlighted. The intensity levels are capped at 100%. See Fig. S7E for the full plot. **(F) ATG8A-IDR binding involves sAIM1-2 and the cAIM regions.** NMR chemical shift perturbations (CSP) of AtC53 IDR (100 µM) in the presence of 75 µM (blue), 100 µM (green), 200 µM (orange) and 300 µM (red) ATG8A. **(G) AtC53 IDR spectra signals in the binding sites shift and broadened upon ATG8 addition.** Insets of overlaid ^1^H-^15^N HSQC spectra of isotope-labeled AtC53 IDR (100 µM) showing chemical shift perturbations of individual peaks from backbone amides of AIM residues in their free (gray) or bound state to unlabeled ATG8A. Unassigned AtC53 IDR residues are indicated by hashtags and HN resonances for residues that could not be assigned in the bound state are shown as gray bars (showing intensity signals of neighbor signals). Chemical shifts are indicated with arrows. Titrations with different concentrations of the ligands are colored similarly to C and F.

Following the backbone assignment, we mapped UFM1 and ATG8 interaction sites in AtC53 IDR by acquiring 2D HSQC spectra of ^15^N-labelled AtC53 IDR in the presence and absence of unlabelled AtUFM1 or ATG8A. Upon AtUFM1 binding, the signals of AtC53 IDR displayed both chemical shift perturbations (CSP) and reduction in their intensity. CSP analysis showed that upon AtUFM1 binding, the signals corresponding to Asp275, Thr279 (sAIM1), Asp286 and Ser297 (sAIM2) and the residues Glu281 and Glu285 that are located between sAIM1 and sAIM2 shifted in a concentration dependent manner (Fig. S7B). Instead, the signals corresponding to Ile274 and Trp276 found in sAIM1, Ile278 and Val280 found in the region between sAIM1 and sAIM2 and Trp287 located in sAIM2 exhibited line broadening and reduced intensity upon binding of AtUFM1 (Fig. 4B). These data confirm that sAIM1 and sAIM2 regions are the major interaction sites for AtUFM1. Notably, the hydrophobic residues between these sAIMs also contributed to the binding. The sAIM1 region showed a significant decrease in signal intensity already at the lowest UFM1 concentration, confirming sAIM1 is the highest affinity binding site for UFM1, followed by sAIM2 region (Fig. 4B-D, S7C). These results are in line with the pulldown assays performed with the Trp to Ala mutants of the sAIMs (Fig. 2H).

We next characterized the binding of ATG8A to AtC53 IDR. Upon ATG8A binding, large number of signals in the AtC53 IDR spectrum disappeared or shifted (Fig. 4E-F, Fig. S7D). The signals of the cAIM and its neighbouring residues covering Leu301 to Glu314 disappeared or shifted at lowest ATG8A concentration (75 µM), followed by sAIM1 and sAIM2 regions as we titrated increased concentrations of ATG8A (Fig. S7E, Fig. 4E). Importantly, the signals Ile274 and Trp276 in sAIM1, which disappeared upon 75 µM UFM1 titration, only disappeared upon 200 µM ATG8A addition, suggesting that while the most preferred binding site for UFM1 is sAIM1, it is cAIM for ATG8A. Similar to UFM1, CSP analysis showed that the signals in sAIM3 region only shifted at highest ATG8A concentration (300 µM) and did not show significant signal intensity reduction, suggesting sAIM3 is a low affinity binding site for both ATG8A and UFM1 (Fig. 4F, G, S7E). Strikingly, residues covering amino acids that precede sAIM1 (265-272) and between cAIM and sAIM3 (315-332) experienced at least a 3-fold increase in their signal intensity upon ATG8A titration (Fig. S7E). However, they displayed minor chemical shift perturbations, suggesting these residues do not directly bind ATG8A, but their dynamics change upon ATG8A binding. Altogether, these data suggest that certain regions in AtC53 IDR might be found in a conformational ensemble that is modulated upon binding of ATG8 but not UFM1. Also, in contrast to UFM1 binding, ATG8A binding triggers a conformational change in C53 IDR. In sum, although both UFM1 and ATG8 bind the sAIMs, their binding modes are different.

To reveal the binding mode of C53 IDR to UFM1 and ATG8, we next set out to map the binding site of C53 IDR on UFM1 and ATG8 using NMR spectroscopy. The backbone amide residues of HsUFM1 and GABARAP have been assigned previously [34, 35]. We successfully transferred 81% of the available backbone spectral assignments for HsUFM1 and 85% for GABARAP to our 2D HSQC spectra, allowing us to characterize the C53 interaction with both UFM1 and ATG8. We then acquired 2D HSQC spectra of ^15^N-labelled UFM1 and ^15^N-labelled ATG8A/GABARAP in the presence and absence of unlabelled C53 IDR. The CSP analysis showed that the signals of Met1, Ser5, Ile8, Lys19, Glu25, Ala31, Lys34, Phe35, Ala36 and Thr67 of HsUFM1 shifted upon HsC53 IDR binding (Fig. S8A-C). Additional residues such as Val32, Glu39, Thr62, Ala63, Gly64 and Asn65 also experienced lower, yet important CSPs indicating a minor contribution of these residues for C53 IDR interaction (Fig. S8C). When we mapped CSPs onto the three-dimensional structure of HsUFM1, we observed a well-defined interaction site on the UFM1 surface covering the α-helix 1 (31-36) and α-helix 2 (62-67), with contributions from residues in β-strand 1 (Ser5, Ile8) and β-strand 2 (Lys19) (Fig. S8D). The AtC53 IDR binding site converges to a region that is involved in the interaction with the UBA5 LIR/UFIM [26], suggesting C53 sAIM interacts with UFM1 in a similar manner to UBA5 LIR/UFIM. To test whether C53 IDR and UBA5 bind UFM1 similarly in plants, we acquired 2D HSQC spectra of ^15^N-labelled AtUFM1 in the presence and absence of unlabelled AtC53 IDR or AtUBA5 LIR/UFIM peptide. Most of the signals that shifted upon AtC53 IDR binding, followed the same trend when AtUFM1 is titrated with AtUBA5 LIR/UFIM, consistent with a conserved binding mode (Fig. S8E). Furthermore, mutation of the tryptophan residue in sAIM1 (AtC53 IDR^W276A^) reduced chemical shift perturbations in AtUFM1 spectrum, supporting its dominant role in AtUFM1 binding (Fig. 4C, D, S8E).

We next analysed the HsC53 IDR-GABARAP interaction. The CSP analysis indicated GABARAP residues Tyr25, Val33, Glu34, Lys35, Ile41, Asp45, Lys46, Tyr49, Leu50 and Phe60 formed intermolecular contacts with C53 IDR (Fig. S9A-C). Additional residues such as Lys20, Ile21, Lys23, Ile32, Asp54, Phe62 and Ile64 displayed smaller CSPs indicating a minor contribution of these residues in the interaction (Fig. S9C). Mapping of CSPs onto the three-dimensional structure of GABARAP highlighted the well-defined LIR docking site (LDS) on the GABARAP surface (Fig. S9D), composed of α-helix 2 (20-25), β-strand 2 (49-52) and α-helix 3 (56-68) residues. Canonical LIR/AIM binding involves the formation of an intermolecular β-sheet with β-strand 2 on ATG8-family proteins and the accommodation of the aromatic and aliphatic residues on two hydrophobic pockets (HP): HP1, which comprises residues in α-helix 2 and β-strand 2, and HP2, formed between the β-strand 2 and α-helix 3, commonly referred to as W and L-site, respectively [36]. However, C53 IDR binding to GABARAP also induces CSPs for residues in β-strand 1 (28-35), closed to α-helix 1 (Fig. S9D). This region has been reported to undergo conformational changes that leads to the formation of a new hydrophobic pocket (HP0) in GABARAP surface upon HsUBA5 LIR/UFIM binding [27]. This suggests, like UFM1, C53 sAIM-ATG8 binding mechanism is similar to UBA5 LIR/UFIM. We confirmed that these binding features are also conserved in plants by acquiring the 2D HSQC spectra of ^15^N-labelled ATG8A in the presence and absence of unlabelled AtC53 IDR or AtUBA5 LIR/UFIM peptide. As for UFM1, most of the signals that shifted followed the same trend upon titration with either C53 IDR or UBA5 LIR/UFIM, demonstrating both motifs bind to a similar site on ATG8 (Fig. S9E). However, unlike UFM1, mutating the aromatic residue in sAIM1 (AtC53 IDR^W276A^) did not reduce CSPs in ATG8A spectrum (Fig. S9D), since binding can proceed via sAIM2 and cAIM residues.

### C53 sAIMs are crucial for C53-mediated autophagy and ER stress tolerance

Our evolutionary and structural analyses suggest that the sAIMs evolved and were selected for their ability to interact with both UFM1 and ATG8. What would happen if we converted sAIMs to cAIMs? We hypothesized that converting sAIMs into cAIMs would reduce the affinity of C53 towards UFM1 and lead to C53 autoactivation, even in the absence of ER stress (Fig. 5A). To test this hypothesis, we generated an AtC53^cAIM^ mutant by re-ordering the residues of each sAIM from IDWD to WDDI. We first assessed the interaction of AtC53^cAIM^ with ATG8A by *in vitro* pulldowns. AtC53^cAIM^ bound ATG8A stronger than the wild type C53 protein. Like the wild type C53 protein, AtC53^cAIM^ interacted via the LIR Docking Site (LDS), as observed by competition with cAIM peptide and loss of interaction in the ATG8^LDS^ mutant (Fig. 5B). On the other hand, AtC53^cAIM^ almost completely lost its ability to bind UFM1, consistent with the dependence of UFM1-binding on the sAIMs (Fig. 5C).

**Figure 5.**
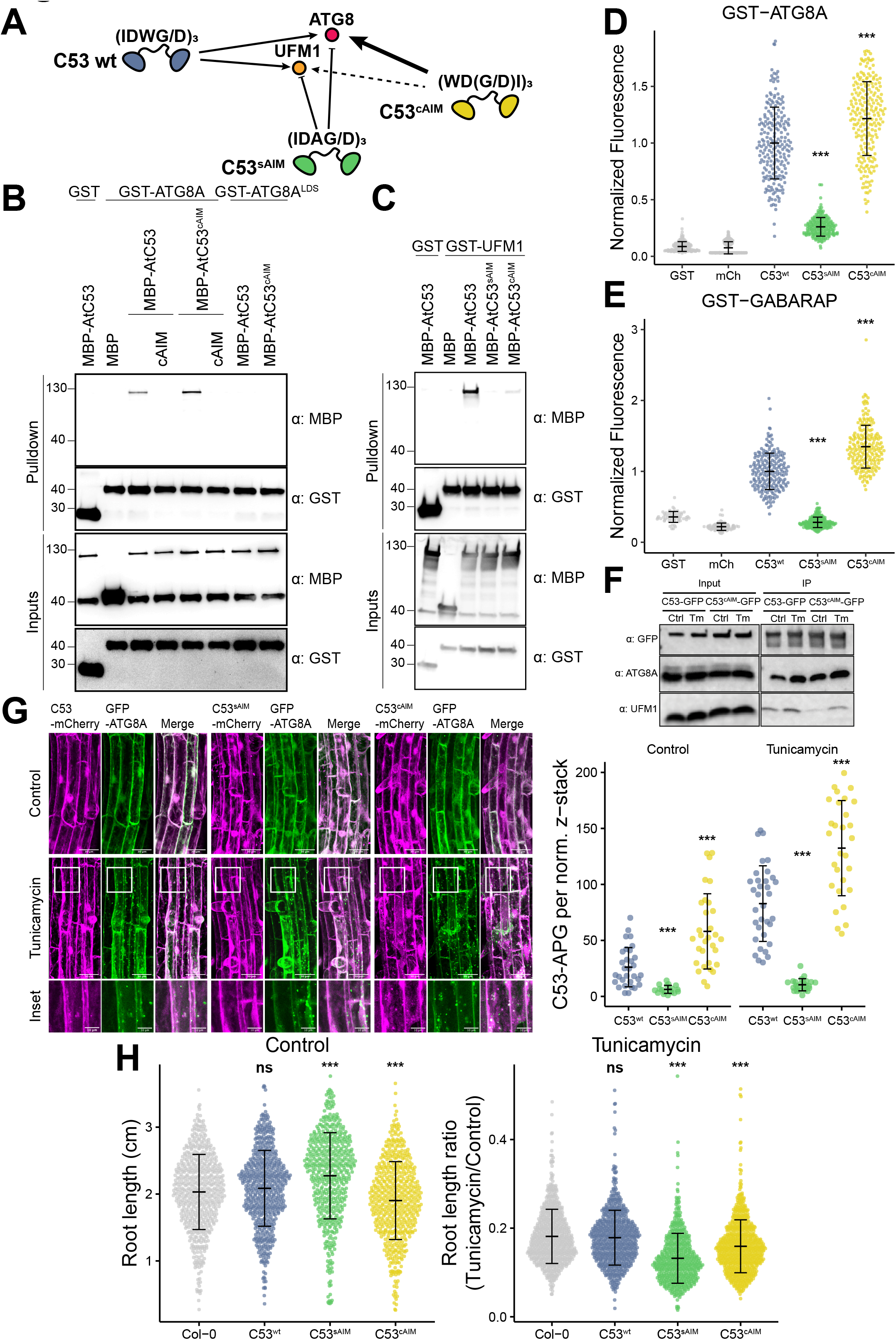
C53 sAIM sequences are essential for ER stress tolerance. **(A) Diagram summarizing our hypothesis that conversion of sAIMs to cAIMs would prevent C53-UFM1 interaction and strengthen C53-ATG8 interaction. (B, C) Conversion of sAIM into cAIM leads to reduced UFM1 binding and stronger ATG8 interaction.** Bacterial lysates containing recombinant proteins were mixed and pulled down with glutathione magnetic agarose beads. Input and bound proteins were visualized by immunoblotting with anti-GST and anti-MBP antibodies. AtC53^sAIM^: AtC53 ^(W276A, W287A, W335A)^; AtC53^cAIM^: AtC53^(IDWD274WDDI, IDWD285WDDI, IDWD333WDDI)^_;_ ATG8^LDS^: ATG8^YL50AA^. **(D, E) Microscopy-based protein–protein interaction assays showing C53^cAIM^ has increased affinity towards ATG8 or GABARAP.** Glutathione-sepharose beads were prepared by incubating them with GST-ATG8A (D) or GST-GABARAP (E). The pre-assembled beads were then washed and mixed with (D) 1 µM of HsC53, 1 µM of HsC53^sAIM^ or 1 µM of HsC53^cAIM^ mutants or (E) 1 µM of AtC53, 1 µM of AtC53^sAIM^ or 1 µM of AtC53^cAIM^ mutants. HsC53^sAIM^: HsC53^(W269A, W294A, W312A)^; HsC53^cAIM^: HsC53^(IDWG267WDGI, IDWG292WDGI, IDWG310WDGI)^. The beads were then imaged using a confocal microscope. Representative confocal images for each condition are shown in figure S10A, B. Normalized fluorescence is shown for each condition with the mean (± SD) of 3 replicate. Unpaired two-samples Wilcoxon test with continuity correction was performed to analyze the differences between wild type and mutants. ***, p-value < 0.001. Total number of beads, mean, median, standard deviation and p-values are reported in Supplementary data 2. **(F) *In vivo* pull downs showing sAIM to cAIM conversion strengthens C53-ATG8 association and weakens C53-UFM1 association.** 6-day old *Arabidopsis* seedlings expressing AtC53-GFP, AtC53^cAIM^-GFP in *c53* mutant background were incubated in liquid 1/2 MS medium with 1% sucrose supplemented with DMSO as control (Ctrl) or 10 μg/ml tunicamycin (Tm) for 16 hours and used for co-immunoprecipitation. Lysates were incubated with GFP-Trap Magnetic Agarose, input and bound proteins were detected by immunoblotting using the respective antibodies as indicated. **(G) AtC53^cAIM^ forms more GFP-ATG8A colocalizing puncta upon ER stress.** *Upper Panel*, representative confocal images of transgenic *Arabidopsis* seedlings co-expressing C53-mCherry (magenta), C53^sAIM^-mCherry and C53^cAIM^-mCherry with GFP-ATG8a in *c53* mutant background under normal condition and after tunicamycin stress. 6-day old seedlings were incubated in liquid 1/2 MS medium with 1% sucrose supplemented with DMSO as control or tunicamycin (10 μg/ml) for 6 hours before imaging. Scale bars, 30 μm. Inset scale bars, 10 μm. *Right Panel*, Quantification of the C53-autophagosomes (C53-APG) per normalized Z-stacks. Bars represent the mean (± SD) of at least twenty roots from 3 biological replicates for each genotype and treatment. Unpaired two-samples Wilcoxon test with continuity correction was performed to analyze the differences between wild type and mutants. ***, p-value < 0.001. **(H) AtC53^cAIM^ mutant is sensitive to ER stress.** Root length quantification of 7-day old *Arabidopsis* seedlings grown vertically on sucrose-free 1/2 MS agar plates supplemented with DMSO control (*Left Panel*, absolute root length in centimeters (cm)) or 100 ng/ml tunicamycin (*Right Panel*, ratio between the root length of tunicamycin treated seedlings and the average of respective control condition). T4 transgenic lines expressing C53-GFP, C53^sAIM^-GFP and C53^cAIM^-GFP in *c53* mutant background were used. Statistical results of more than 500 seedlings from 3 biological repeats per each genotype for control and tunicamycin treated condition are shown. Bars represent the mean (± SD) of 3 biological replicates. Unpaired two-samples Wilcoxon test with continuity correction was performed to analyze the differences between wild type and mutants. ns, p-value > 0.05, ***, p-value < 0.001.

To further corroborate our *in vitro* pulldown assays, we performed quantitative on-bead binding assays. GST-ATG8 and GST-GABARAP recruited C53^cAIM^ mutant 22% (mean) and 35% (mean) more efficiently than the respective C53 wild type proteins (Fig. 5D, 5E, S10A, S10B). C53^sAIM^ mutant (with inactivated sAIMs) was instead recruited 74% (mean) and 78% (mean) less to GST-ATG8 and GST-GABARAP, respectively (Fig. 5D, 5E, S10A, S10B). In addition to ATG8, C53 also interacts with the scaffold protein FIP200/ATG11 [37, 38]. We therefore tested the binding affinities of C53 and C53^cAIM^ to FIP200. Similar to our observations with ATG8, HsC53^cAIM^ displayed a stronger interaction with FIP200 than wild type HsC53. Similar to ATG8, FIP200 interaction was also lost in C53^sAIM^ mutant (Fig. S11). These results demonstrate that converting sAIM to cAIM increases the affinity of C53 towards ATG8 and decreases its affinity to UFM1.

We next explored the physiological consequences of sAIM to cAIM conversion. We complemented an *Arabidopsis thaliana c53* mutants with either C53-GFP, C53^sAIM^-GFP, or C53^cAIM^-GFP fusions. Consistent with our *in vitro* data, *in vivo* pull-down assays showed that C53^cAIM^-GFP had a stronger interaction with ATG8 than C53-GFP. On the contrary, the association between C53^cAIM^-GFP and UFM1 was weaker than between C53-GFP and UFM1 (Fig. 5F, S12A, S12B).

Under normal conditions, Arabidopsis C53 predominantly has a diffuse cytoplasmic localization pattern. Upon ER stress, it is recruited to the ATG8-labelled autophagosomes [14]. Consistent with our *in vivo* pull-down results, C53^cAIM^-mCherry formed puncta even under normal conditions, suggesting it associates with ATG8 and recruited to the autophagosomes even in the absence of stress. Altogether, these findings suggest sAIM to cAIM conversion leads to the premature activation of C53-mediated autophagy (Fig. 5G).

Finally, using tunicamycin plate assays, we measured ER stress tolerance of C53^cAIM^ expressing Arabidopsis plants. Tunicamycin is a glycosylation inhibitor that is commonly used to induce ER stress in plants, which leads to the shortening of the roots in *Arabidopsis thaliana* [39]. Compared to wild type complemented plants, C53^cAIM^ expressing Arabidopsis lines formed shorter roots even under control conditions (Fig. 5H). This suggests, premature activation of C53 is detrimental for plant growth, likely due to the degradation of C53 without the bound cargo. The root length was further reduced in tunicamycin containing plates, indicating the inability to degrade C53 cargo that arise upon ER stress is detrimental for plants. Taken together, our results illustrate that C53’s ability to bind UFM1 and ATG8, which is encoded in sAIM regions, is crucial for its function and ER stress tolerance.

## Discussion

Despite the discovery of UFMylation almost two decades ago, its structural basis, the full spectrum of UFMylated substrates, and its physiological role are still not fully resolved [17, 40]. Studies in metazoans and our recent work have shown that UFMylation is involved in a wide range of homeostatic pathways, including ER stress tolerance, immunity, autophagy, lipid droplet biogenesis, and the DNA damage responses [14, 15, 23, 41–46]. In ER homeostasis, UFMylation is activated by stalling of ER-bound ribosomes and brings about the degradation of incomplete polypeptides, which can be toxic for the cell [14, 23]. Limited phylogenetic analysis, comparing yeast to plants and metazoans, suggested that the pathway had evolved in multicellular eukaryotes and could have facilitated the protein synthesis burden that arises during biogenesis of the extracellular matrix [22]. However, our extensive phylogenomic analysis, in agreement with a recent study, clearly shows that UFMylation did not evolve in multicellular eukaryotes, but was secondarily lost in fungi and other lineages [47] (Fig. 1). Indeed, many single-celled organisms including *Chlamydomonas* harbour a full complement of UFMylation components in their genome, whereas certain multicellular lineages, such as kelp (Phaeophyceae), have lost the majority of the pathway. We provide biochemical and physiological evidence showing UFMylation is functional in *Chlamydomonas*, unequivocally refuting the idea that UFMylation evolved only in multicellular organisms (Fig. 1). Our evolutionary analysis also highlights why we should move beyond yeast and metazoans and instead consider the whole tree of life when using evolutionary arguments to guide biological research. Our phylogenetic analysis also revealed that in addition to the Fungi, several algal groups, and pathogens such as *Plasmodium*, *Entamoeba,* and *Trichomonas* have also lost UFMylation. So, how do pathogens and parasitic fungi resolve stalled ER-bound ribosomes? Comparative studies addressing these questions could provide potential translational avenues for developing genetic or chemical means to prevent infections.

Another conclusion of our phylogenetic studies is the tight connection between the presence of sAIMs located in the C53 IDR and UFM1. Species that lack UFM1 also lost the sAIMs in C53 (Fig. 2). Using biochemical and structural approaches, we found that sAIMs form versatile docking sites that can interact with both UFM1 and ATG8. UFM1 interaction is mostly mediated by sAIM1 and sAIM2, whereas ATG8 interaction is driven by the cAIM, sAIM1 and sAIM2 (Fig. 2, Fig. 4). It is surprising that the sAIM3, which is highly similar to sAIM1/2 does not show significant binding to UFM1. A plausible explanation is that the aspartic acid at the second position in sAIM1/2 (I**D**WD) motif play an important role for the interaction, and having a serine instead of an aspartic acid in sAIM3 (I**S**WD) weakens the binding. Consistently, the NMR analyses showed that the signals of the residues neighboring sAIMs showed significant chemical shifts suggesting that they also contribute to the interaction with both UFM1 and ATG8.

The NMR experiments also revealed that UFM1 and ATG8 binding induce distinct conformational changes on C53 IDR (Fig. S7). UFM1 binding reduces the overall signal intensity with further reduction at the direct binding sites corresponding to sAIM1 and sAIM2. On the contrary, ATG8 binding leads to a local signal intensity drop at the sAIM1-2 and cAIM but increases the signal intensity of residues that do not interact with ATG8. These data suggest that upon ATG8 binding C53 IDR becomes more dynamic, potentially allowing it to bind the autophagic cargo. This structural rearrangement could also affect the E3 ligase activity of the UFL1 enzyme complex. Indeed, a recent study has shown that C53 negatively regulates UFMylation activity, when bound to the UFL1-DDRGK1 complex [21]. Altogether, these results indicate that evolution of suboptimal ATG8 interacting motifs enabled C53 to interact with another regulatory protein, UFM1, creating an autoinhibition mechanism that regulates ER-phagy. This illustrates how complex regulatory circuits could evolve by shuffling existing short linear motifs.

Interestingly, another non-canonical motif on UBA5, the E1 enzyme of the UFMylation cascade, can also bind both UFM1 and ATG8 through similar binding pockets (Fig. S8, Fig. S9). Removing or mutating UBA5 LIR affects the kinetics of UFMylation and the GABARAP dependent recruitment of UBA5 to ER upon stress [27]. Our findings go a step further and show that non-canonical motifs on C53 are essential for organismal fitness, as converting sAIMs to canonical AIMs leads to reduced ER stress tolerance in *Arabidopsis thaliana* (Fig. 5H). Further *in vitro* reconstitution studies that involve the UFMylation machinery, C53 receptor complex, and stalled membrane-bound ribosomes are necessary to understand the dynamic changes that lead to C53 activation, which would explain how UFMylation and autophagy intersect at the ER.

In summary, our data converge on the model that UFM1 and ATG8 compete for C53 binding via the shuffled ATG8 interacting motifs [14]. Under normal conditions, C53 is bound to UFM1, keeping it inactive. Upon stress, UFM1 is displaced by ATG8, leading to structural rearrangements that trigger C53-mediated autophagy. These results provide a mechanism where the cell keeps selective autophagy pathways inactive under normal conditions to prevent the spurious degradation of healthy cellular components and saves the energy that is required to form autophagosomes.

## Acknowledgements

We thank Vienna Biocenter Core Facilities (VBCF) Protein Chemistry, Biooptics, Plant Sciences, Molecular Biology, and Protein Technologies. We acknowledge funding from Austrian Academy of Sciences, Austrian Science Fund (FWF, P 32355, P 34944), Austrian Science Fund (FWF-SFB F79), Vienna Science and Technology Fund (WWTF, LS17-047), Marie Curie Fellowship to NZ. RB acknowledges support of a UKRI Future Leaders Fellowship (Grant Reference MR/T020970/1) and a Chancellor’s Fellowship from the University of Strathclyde. The IMP is supported by Boehringer Ingelheim. We thank J. Matthew Watson and members of the Dagdas lab for the critical reading and editing of the manuscript. We are thankful to Max Perutz Labs NMR facility and Georg Kontaxis for the NMR measurements. We thank Mathias Madalinski, Zsuzsanna Muhari-Portik for peptide synthesis. We also thank Orla Dunne, Arthur Sedivy, David Drechsel for help in protein purification and characterization experiments.

## Contributions

LP, VM, NZ, EK and YD conceived and designed the project. NI performed the phylogenomic analysis. LP and SR performed the *Chlamydomonas reinhardtii* related experiments. LP, VM and JSM performed *in vitro* biochemical and biophysical assays. RB performed native mass spectrometry experiments. TL trained the deep learning model for agarose bead recognition. VM and HH performed the NMR spectroscopy experiments. NZ, MS and NG performed *Arabidopsis thaliana* related experiments. SM, TR, TC, SR, EK and YD supervised the project. LP, VM, NZ, NI, EK and YD wrote the manuscript with input from all the authors.

## Materials and Methods

### Phylogenomic analysis

To reconstruct the evolutionary history of the UFMylation pathway, we searched for UFMylation proteins (including RPL26) in 151 eukaryotic datasets comprising 149 genomes and two transcriptomes from the dinoflagellates *Togula jolla* and *Polarella glacialis* (Supplementary Data S1). Initially, *Homo sapiens* proteins were used as queries to search predicted proteomes using Diamond BLASTp v2.0.9 (E-value < 10^-5^, ultra-sensitive mode) [48]. Multiple sequence alignments were then inferred using MAFFT v7.490 (-auto) and trimmed using trimAl v1.4 with a gap-threshold of 30%, before preliminary phylogenies were generated using IQ-Tree v2.1.2 (LG4X model, fast mode) [49–51]. The resulting phylogenies were annotated using SWISS-PROT (version 2022_01) and Pfam (version 35.0) and then interpreted in FigTree v1.4.2. From the phylogeny, orthologs were identified, extracted, and used as queries for a second iteration of BLAST searching as described above [52–54]. To improve search sensitivity, the orthologs identified using BLAST were then used to generate profile hidden Markov models (HMMs). Initially, the proteins were re-aligned with the structurally informed aligner MAFFT-DASH with the L-INS-i algorithm and were then trimmed with a gap-threshold of 10% [55]. HMMs were then generated from the alignments and used to re-search the proteomic datasets using HMMER v3.1b2 (E-value < 10^-5^) [56]. The identified homologs were once again aligned, trimmed, and assessed phylogenetically, facilitating the removal of paralogs. Lastly, to account for the possibility that proteins could be missing due to genomic mis-annotation, proteins identified from the predicted proteomes were used as queries for tBLASTn (E-value < 10^-5^) searches against eukaryotic genomes and protein predictions were generated using Exonerate v2.2 (see https://github.com/nickatirwin/Phylogenomic-analysis) [57]. Newly predicted proteins were combined with the previously identified proteins and were once again phylogenetically screened for paralogs. The presence and absence of the resulting orthologs was plotted across a eukaryotic phylogeny using ITOL v6 with taxonomic information inferred from NCBI Taxonomy following adjustments made based on recent phylogenomic analyses [58–60].

To investigate the sequence conservation of C53 and RPL26, multiple sequence alignments were generated from the identified orthologs using MAFFT with the L-INS-i algorithm. The alignments were then trimmed using a gap-threshold of 30% and fragmented sequences with less than 50% data were filtered out. In the case of C53, alignment of the poorly conserved intrinsically disordered region (IDR) was improved through re-alignment using MUSCLE v3.8 implemented in AliView v1.26 [61, 62]. For C53, phylogenetic analyses were conducted using IQ-Tree and substitution models were selected using ModelFinder (LG+F+R6) [63]. The phylogeny and C53 alignment were then used in an analysis using ConSurf to examine sequence conservation. Likewise, both the C53 and RPL26 alignments were used to assess sequence conservation and divergence between species with and without UFM1 using TwinCons (using the LG substitution model and Voronoi clustering) [64]. Lastly, alignment logos for the C53 shuffled AIMs were generated with Skylign using weighted counts [65].

### Cloning procedures

Constructs for *Arabidopsis thaliana* and *Escherichia coli* transformation were generated using the GreenGate (GG) cloning method [66]. Plasmids used are listed in materials section. The coding sequence of genes of interest were either ordered from Twist Biosciences or Genewiz or amplified from Col-0 using the primers listed in the materials section. The internal *Bsa*I sites were mutated by site-directed-mutagenesis without affecting the amino acid sequence.

### Chlamydomonas reinhardtii genomic DNA extraction

The following protocol was adapted from *Perlaza K., et al. 2019* [67]. A 6 ml aliquot of a liquid TAP culture in mid-log phase was spun down, and the media was decanted. The pellet was resuspended in 400 µl of water and then 1 volume of DNA lysis buffer was added (200 mM Tris HCl pH 8.0, 6% SDS, 2 mM (EDTA)). To digest proteins, 5 µ of 20 mg/ml proteinase K (Thermo Fischer) was added and allowed to incubate at Room Temperature (RT) for 15 min. 200 µl of 5M NaCl was then added and mixed gently. Next, to selectively precipitate nucleic acids, 160 µl of 10% CTAB in 0.7 M NaCl was added and allowed to sit for 10 min at 65°C with gentle agitation. Two or more consecutive rounds of DNA extraction using ultrapure phenol:chloroform:isoamyl alcohol (25:24:1, v/v/v) were performed to achieve a clean interphase. Then, the upper aqueous phase was retained and mixed with 1 volume of 2-propanol. This was mixed gently for 15 min at RT. Then it was spun down for 30 min at 21,000 x g at 4°C. The supernatant was removed and 1 volume of ice-cold 70% ethanol was added and mixed with the pellet. This mixture was spun down for 15 min at 21,000 x g. The supernatant was removed, and the DNA precipitate was dried in a speed-vac for about 10– 25 min and resuspended in 40 µl of nuclease-free water.

The purity of the genomic DNA preparation was assessed using a spectrophotometer, ensuring absorbance ratios at 260/280 nm and 260/230 nm to be ∼1.8 and ∼2.0, respectively, prior to using the genomic DNA preparation for most of the follow-up applications.

### Genotyping of the Chlamydomonas reinhardtii mutants

The insertion of the mutagenic cassette (PARO) in the UBA5 and UFL1 loci was verified by PCR by using primers designed to anneal inside and outside of the PARO cassette, using KOD Extreme Hot Start DNA Polymerase (Sigma). The PCR products were run on 1 % (w/v) agarose. The primer sequences and expected PCR products can be found in *Materials*.

### Chlamydomonas reinhardtii in vivo UFMylation assays

Cell cultures were grown in liquid TAP medium in 100 ml Erlenmeyer flasks for about two days to an OD_600_ of 1.5-2. These cultures were then transferred to fresh liquid TAP medium, with or without 0.2 mg/l Tunicamycin, to a final OD_600_ of 0.1. After either 12 hours or 24 hours of treatment, 5 ml of cell culture was spun down, flash frozen in liquid nitrogen and stored at −70 °C.

The pellets were thawed and resuspended in 150 µl of SDS-lysis buffer (100 mM Tris-HCl pH 8.0, 600 mM NaCl, 4% SDS, 20 mM EDTA, freshly supplied with Roche Protease Inhibitors). Samples were vortexed for 10 min at RT and centrifuged at maximum speed for 15 min at 4°C to remove the cell debris. The supernatant, containing a total extract of denatured proteins was transferred to a new eppendorf tube, a 5 µl aliquot was saved for BCA quantification and diluted accordingly.

5X SDS-loading buffer (250 mM Tris-HCl pH 6.8, 5% SDS, 0.025% bromophenol blue, 25% glycerol), freshly supplied with 5% of β-mercaptoethanol, was added to the extract and denatured at 90°C for 10 min. The samples were loaded on 4–20% SDS-PAGE gradient gel (BioRad) and electrophoresis was run at 100V for 1.5 hr.

### Chlamydomonas reinhardtii survival assays

Cell cultures were grown in liquid TAP medium in a 100 ml Erlenmeyer flask for about two days to an OD_600_ of 1.5-2. These cultures were then transferred to fresh liquid TAP medium, with or without 0.2 mg/l Tunicamycin, to a final OD_600_ of 0.1. After 24, 48 and 72 hours of treatment, the optical density (OD) of the cultures was measured using a spectrophotometer at 600 nm.

### *Arabidopsis thaliana* plant materials and growth conditions

The Columbia-0 (Col-0) accession of *Arabidopsis* was used in this study unless otherwise indicated. *Arabidopsis* mutants used in this study are listed in the materials section. Generation of transgenic *Arabidopsis* plants was carried out by *Agrobacterium*-mediated transformation [68].

Seeds were imbibed at 4°C for 3 days in dark. For the co-immunoprecipitation experiment, seeds were sterilized and cultured in liquid 1/2 MS medium containing 1% sucrose with constant shaking under continuous LED light. For the root length measurements, seeds are sterilized and sown on sucrose-free 1/2 MS agar plates and grown at 22°C at 60% humidity under continuous white light at 12/12-hour light/dark cycle.

### Root length quantification

Seedlings were grown vertically for 7 days on sucrose-free 1/2 MS plates supplemented with indicated chemicals. Plates were photographed using a Canon EOS 80D camera. The root length was measured using ImageJ software (version: 2.1.0/1.53c) for further analysis [69].

### *In vivo* co-immunoprecipitation

*Arabidopsis* seedlings were cultured in liquid 1/2 MS medium with 1% sucrose for 7-8 days. These seedlings were then treated for additional 16 hours in 1/2 MS liquid medium with 1% sucrose supplemented with DMSO or tunicamycin, respectively. About 1-2 mg plant material was harvested and homogenized using liquid nitrogen and immediately dissolved in grinding buffer (50 mM Tris-HCl, pH 7.5, 150 mM NaCl, 10 mM MgCl_2_, 10% glycerol, 0.1% Nonidet P-40, Protease Inhibitor Cocktail tablet) by vortex. Plant lysates were cleared by centrifugation at 16,000g for 5 min at 4°C several times. After binding to Protein A Agarose, 3 mg total plant protein were incubated with 25 µL GFP-Trap Magnetic Agarose beads (ChromoTek) at 4°C for 2.5 hours. Pellets were washed with grinding buffer for six times, boiled for 10 min at 95°C prior to immunoblotting with the respective antibodies.

### Confocal microscopy

*Arabidopsis* roots were imaged using a Zeiss LSM780 confocal microscope with an Apochromat 20x objective lens at 2 X magnification. Z-stack merged images with 2 μm thickness per Z-stack were used for analysis. At least 5 Z-stacks were used for puncta quantification and image presentation. Confocal images were processed with ImageJ software [69].

### Quantification of confocal micrographs

ImageJ software (version: 2.1.0/1.53c) [69] is used for autophagic puncta number quantification. ATG8A puncta colocalized C53 punctuates were manually mounted for each stack and added for all stacks for a single image. Autophagosome number per normalized Z-stack was calculated by total autophagosome number of a certain image divided by the relative root area.

### Western blotting

Blotting on nitrocellulose membranes was performed using a semi-dry Turbo transfer blot system (BioRad). Membranes were blocked with 5% skimmed milk or BSA in TBS and 0.1% Tween 20 (TBS-T) for 1 hour at room temperature or at 4°C overnight. This was followed by incubation with primary and subsequent secondary antibody conjugated to horseradish peroxidase. After five 5 min washes with TBS-T, the immune-reaction was developed using either Pierce™ ECL Western Blotting Substrate (ThermoFisher) or SuperSignal™ West Pico PLUS Chemiluminescent Substrate (ThermoFisher) and detected with either ChemiDoc Touch Imaging System (BioRad) or iBright Imaging System (Invitrogen).

### Western blot image quantification

Protein bands intensities were quantified with ImageJ [69]. Equal rectangles were drawn around the total protein gel lane and the band of interest. The area of the peak in the profile was taken as a measure of the band intensity. The protein band of interest was normalized for the total protein level of the protein lane used as a bait. Average relative intensities and a standard error of three independent experiments were calculated.

### Protein expression and purification for biochemical assays

Recombinant proteins were produced using *E. coli* strain Rosetta2 (DE3) pLysS grown in 2x TY media at 37°C to an A_600_ of 0.4–0.6 followed by induction with 300 µM IPTG and overnight incubation at 18°C.

For *in vitro* UFMylation assays, *in vitro* pulldowns, and *in vitro* protein-protein microscopy binding assays pelleted cells were resuspended in lysis buffer (100 mM HEPES pH 7.5, 300 mM NaCl) containing protease inhibitors (Complete™, Roche) and sonicated. The clarified lysate was first purified by affinity, by using HisTrap FF (GE HealthCare) columns. The proteins were eluted with lysis buffer containing 500 mM imidazole. The eluted fraction was buffer exchanged to 10 mM HEPES pH 7.5, 100 mM NaCl and loaded either on Cation Exchange, Resource S, or Anion Exchange, Resource Q, chromatography columns. The proteins were eluted from 5 to 55 % of Ion exchange buffer B (10 mM HEPES pH 7.5, 1 M NaCl by NaCl) gradient in 20 CV. Finally, the proteins were separated by Size Exclusion Chromatography with HiLoad® 16/600 Superdex® 200 pg or HiLoad® 16/600 Superdex® 75 pg, which were previously equilibrated in 50 mM HEPES pH 7.5, 150 mM NaCl.

The proteins were concentrated using Vivaspin concentrators (3000, 5000, 10000 or 30000 MWCO). Protein concentration was calculated from the UV absorption at 280 nm by DS-11 FX+ Spectrophotometer (DeNovix).

### Protein expression and purification for Nuclear Magnetic Resonance (NMR) spectroscopy

All recombinant proteins were produced using *E. coli* strain Rosetta2 (DE3) pLysS. Transformed cells were grown in 2x TY media supplemented with 100 µg/mL spectinomycin at 37°C to log phase (OD_600_ 0.6-0.8), followed by induction with 300 µM isopropyl β-D-1-thiogalactopyranoside (IPTG) and incubation at 18°C overnight. Recombinant isotopically labelled proteins used for Nuclear Magnetic Resonance (NMR) spectroscopy were grown in M9 minimal media as previously described [70] supplemented in the presence of 100 µg/mL spectinomycin at 37°C to log phase (OD_600_ 0.6-0.8), followed by induction with 600 µM isopropyl β-D-1-thiogalactopyranoside (IPTG) and incubation at 18°C overnight. Cells were harvested by centrifugation and resuspended in lysis buffer of 100 mM Sodium Phosphate (pH 7.0), 300 mM NaCl, 20 mM imidazole supplemented with Complete-EDTA-Free Protease Inhibitor (Roche) and benzonase. Cells were lysed by sonication and lysate was clarified by centrifugation at 20,000 x g. The clarified lysate was loaded on a HisTrapFF (GE Healthcare) column pre-equilibrated with the lysis buffer. Proteins were washed with lysis buffer for 10 CV and eluted with lysis buffer containing 500 mM Imidazole. The eluted fraction was buffer exchanged to 10mM Sodium Phosphate (pH 7.0), 50 mM NaCl and loaded either on Cation Exchange (ResourceS, Cytiva) or Anion Exchange (ResourceQ, Cytiva) chromatography columns. The proteins were eluted by NaCl gradient (50% in 20 CV). Samples were further purified by size-exclusion chromatography with HiLoad 16/600 Superdex 200 pg or HiLoad 16/600 Superdex 75 pg (GE Healthcare) with 50 mM Sodium Phosphate (pH 7.0), 100 mM NaCl. The proteins were concentrated using VivaSpin concentrators (3000, 5000, 10000, or 30000 MWCO). Protein concentration was calculated from the UV absorption at 280 nm by DS-11 FX+ Spectrophotometer (DeNovix) or at 205nm by Jasco V-750 UV-Visible Spectrophotometer.

### *In vitro* UFMylation assays

CrUBA5, CrUFC1 and UFM1 were mixed to a final concentration of 5 µM, 5 µM and 20 µM respectively in a buffer containing 25 mM HEPES pH 7.5, 150 mM NaCl and 10 mM MgCl_2_. The enzymatic reaction was started by adding ATP to a final concentration of 5 µM. The enzymatic mixture was incubated for 1 hour at 37°C and then stopped with the addition of non-reducing Laemmli Loading Buffer. Beta-MercaptoEthanol (BME) was added only where specified to reduce UBA5-UFM1 or UFC1-UFM1 thioester bond. The samples were loaded on 4–20% SDS-PAGE gradient gel (BioRad) and electrophoresis was run at 100V for 1.5 hr.

### *In vitro* pulldowns

For pulldown experiments, 5 µl of glutathione magnetic agarose beads (Pierce Glutathione Magnetic Agarose Beads, Thermo Scientific) were equilibrated by washing them two times with wash buffer (100 mM Sodium Phosphate pH 7.2, 300 mM NaCl, 1 mM DTT, 0.01% (v/v) IGEPAL). Normalized *E. coli* clarified lysates or purified proteins were mixed, according to the experiment, added to the washed beads and incubated on an end-over-end rotator for 1 hour at 4°C. Beads were washed five times with 1 ml wash buffer. Bound proteins were eluted by adding 50 µl Laemmli buffer. Samples were analyzed by western blotting or Coomassie staining.

### Microscopy-based on-bead protein-protein interaction assays

Glutathione Sepharose 4B bead slurry (Cytiva, average diameter 90 µm) was washed and diluted 10 times in HEPES buffer (25 mM HEPES pH 7.5, 150 mM NaCl, 1 mM DTT). The beads were then incubated for 30 min at 4°C (16 rpm horizontal rotation) with GST-tagged bait proteins (2 µM of GST, GST-FIP200 CD, GST-ATG8A, GST-GABARAP, GST-AtUFM1, GST-HsUFM1). The beads were washed 5 times in 10 times the bead volume of HEPES buffer. The buffer was removed, and the beads were resuspended 1:20 in HEPES buffer. 10µl of diluted beads were mixed with 20 µl of mCherry tagged binding partner at a concentration of 1.5 µM (0.5 µl bead slurry and 1 µM binding partner final concentrations) with or without competitor, as stated in the relative experiment. The mixture was transferred to a black, glass bottom, 384-well plate (Greiner Bio-One) and incubated for 30-60 min at RT.

Imaging was performed with either a Zeiss LSM700 confocal microscope with 20 X magnification or with a Zeiss LSM800 confocal microscope with 10 X magnification.

### Quantification of microscopy-based protein-protein interaction assays

From images acquired from a Zeiss LSM700 confocal microscope, the quantification of fluorescence was performed in ImageJ [69] by drawing a line across each bead and taking the maximum gray value along the line. The maximum gray value for any given pixel represents the fluorescence intensity.

For images acquired from a Zeiss LSM800 confocal microscope, we used a custom Fiji Macro. Within this workflow a pretrained model was created for the deep learning application “Stardist” (https://imagej.net/plugins/stardist) [71]. This model was based on a manually annotated training set, using the fluorescently labelled beads as a basis for creating the ground truth annotations, then performing the training on the brightfield channel. Out of focus beads were rejected in this step and therefore excluded from the training. After applying the deep learning-based segmentation, the regions were reduced to a ring around the edge of the beads. Beads on image borders were excluded from the analysis. In the end, the mean fluorescent intensities were exported out and used for quantification.

For each method, the fluorescence intensity was normalized against the mean of the control condition.

Fiji macro and agarose bead model for automatic quantification are available in Supplementary Data 3.

### Mass Spectrometry Measurements

Proteins were buffer exchanged into ammonium acetate using BioRad Micro Bio-Spin 6 Columns. Native mass spectrometry experiments were carried out on a Synapt G2Si instrument (Waters, Manchester, UK) with a nanoelectrospray ionization source (nESI). Mass calibration was performed by a separate infusion of NaI cluster ions. Solutions were ionized from a thin-walled borosilicate glass capillary (i.d. 0.78 mm, o.d. 1.0 mm, Sutter Instrument Co., Novato, CA, USA) pulled in-house to nESI tip with a Flaming/Brown micropipette puller (Sutter Instrument Co., Novato, CA, USA). A potential of 0.8 kV was applied to the solution via a thin platinum wire (diameter 0.125 mm, Goodfellow, Huntingdon, UK). The following instrument parameters were used: capillary voltage 0.8 kV, sample cone voltage 40 V, source offset 60 V, source temperature 40 °C, trap collision energy 4.0 V, trap gas 3 mL/min. Data were processed using Masslynx V4.2 and OriginPro 2021.

### NMR spectroscopy

All NMR spectroscopy measurements were performed using Bruker AVIII 600MHz or Avance 800MHz spectrometers at 25°C. The data were processed using TopSpin 3.2 (Bruker) and NMRPipe [72] and analysed using CcpNmr Analysis [73].

Sequence specific backbone assignments of AtC53 IDR were achieved using 2D ^1^H-^15^N HSQC, 3D HNCA, 3D CBCACONH, 3D HNCACB, 3D HNCO, 3D HNCACO including 70 residues of 75 non-proline residues (93%). NMR titrations were performed by adding unlabelled protein (75-300 µM) to 100 µM of ^15^N single-labelled protein in 50 mM sodium phosphate (pH 7.0), 100 mM NaCl and 10% (v/v) D_2_O and monitored by two-dimensional ^1^H-^15^N HSQC.

### Statistical analysis

All statistical analysis was performed using R Statistical Software (version 4.1.2; R Foundation for Statistical Computing, Vienna, Austria) [74]. Statistical significance of differences between two experimental groups was assessed with a two-tailed unpaired two-samples t-test if the two groups were normally distributed (Shapiro-Wilk test) and their variances were equal (F-test). If the groups were normally distributed but the variances were not equal a two-samples Welch t-test was performed. If the groups were not normally distributed, an unpaired two-samples Wilcoxon test with continuity correction was performed. Differences between two data sets were considered significant at p < 0.05 (*); p < 0.01 (**); p<0.001 (***). P value > 0.05 (ns, not significant).

## MATERIALS

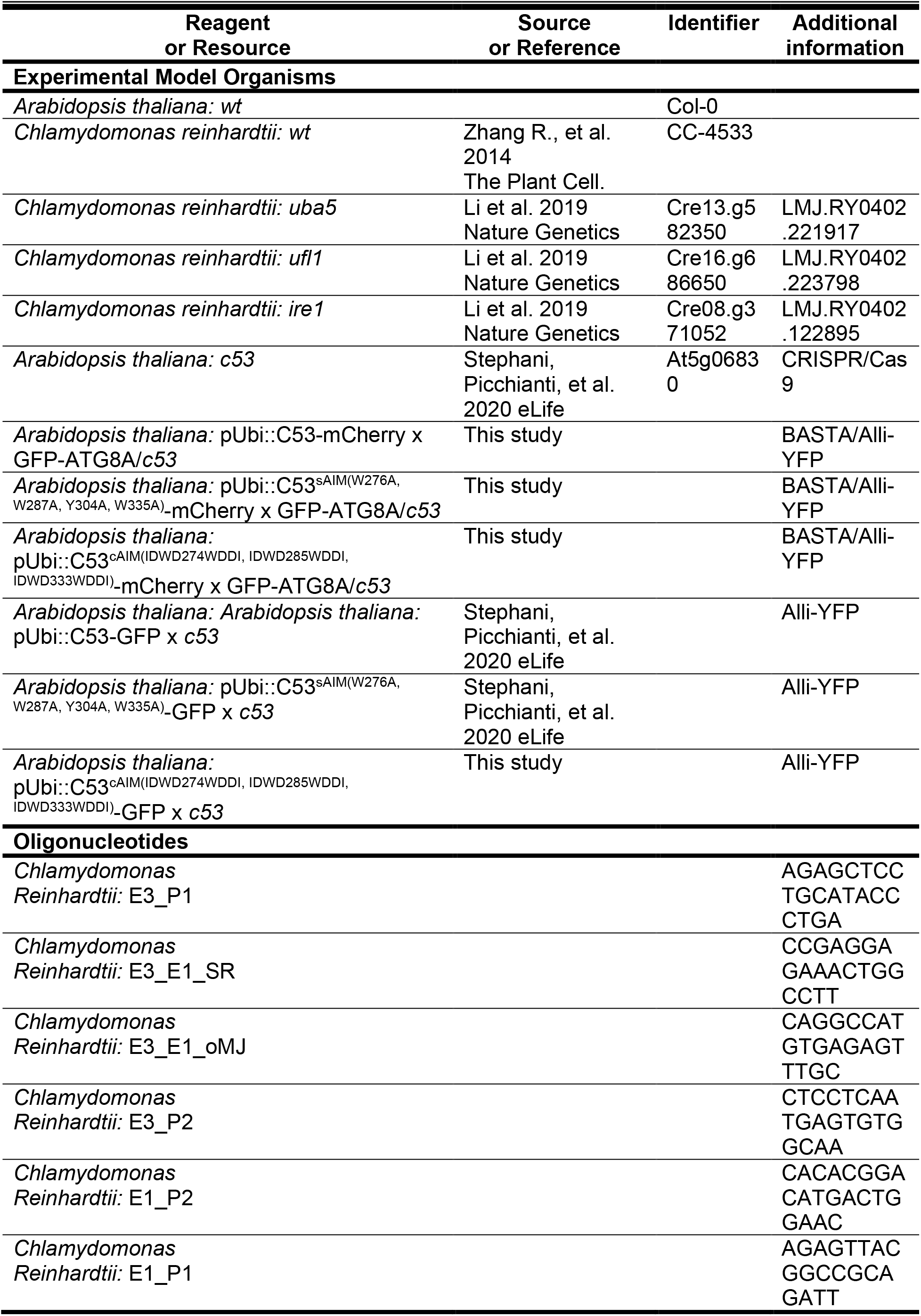
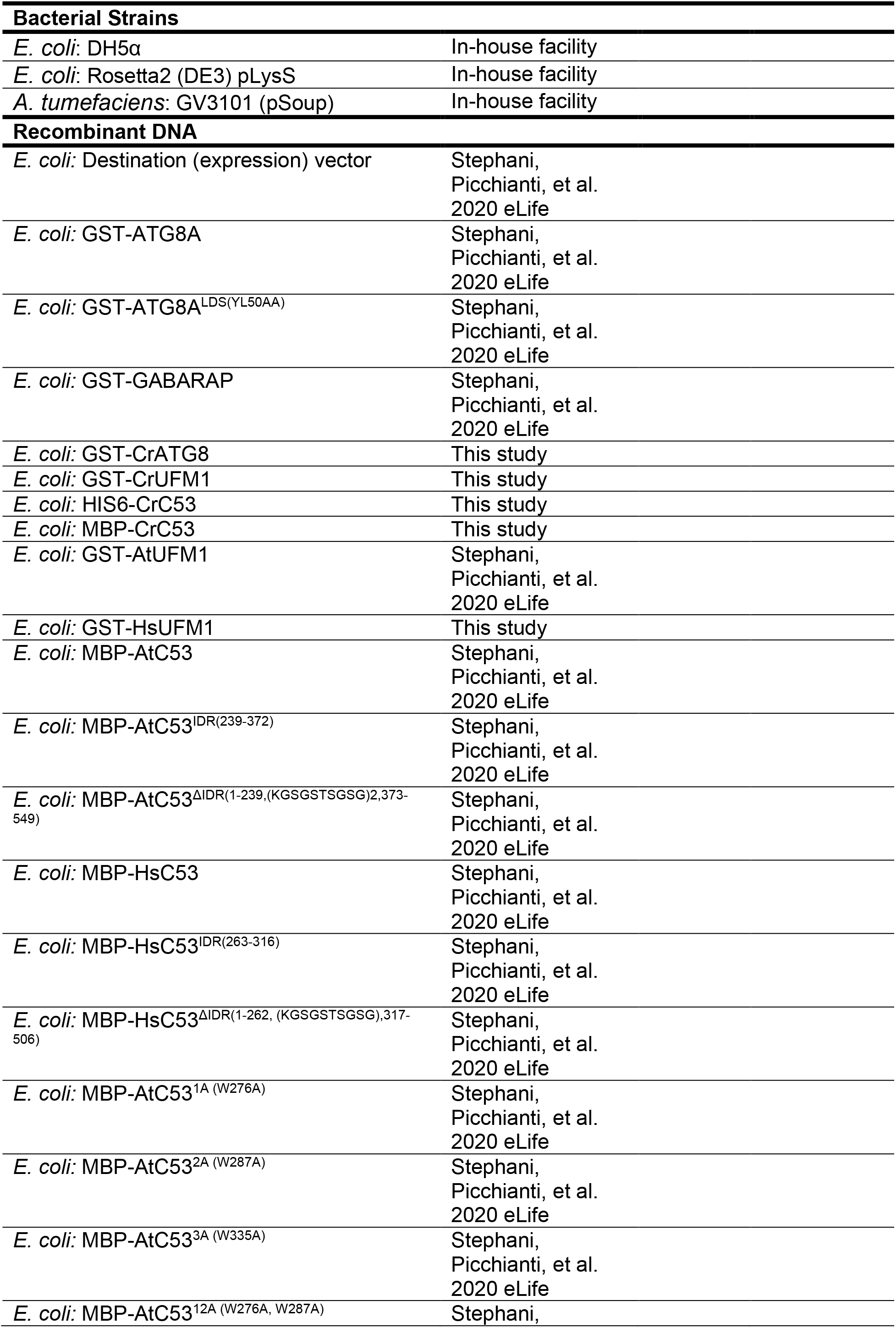
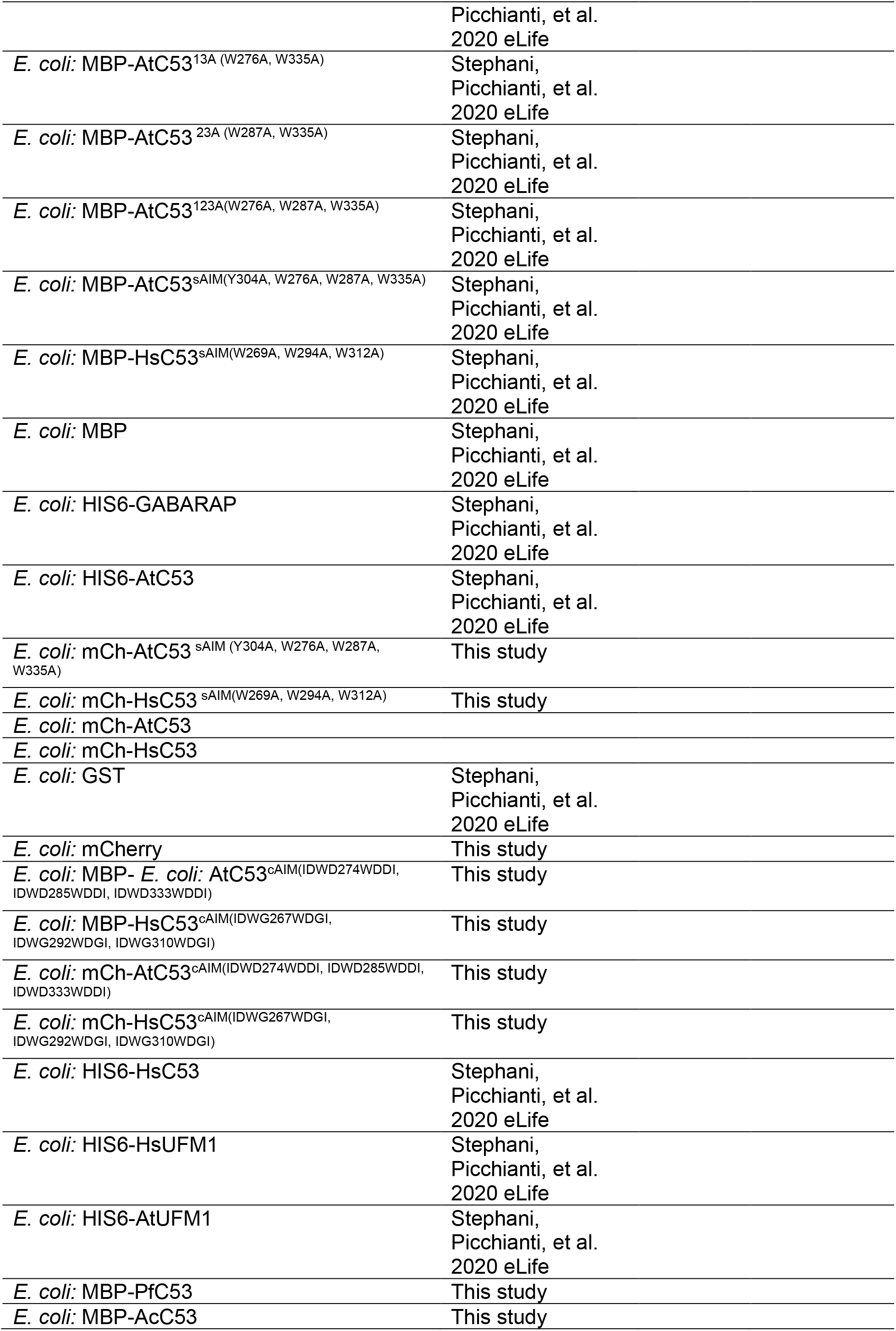
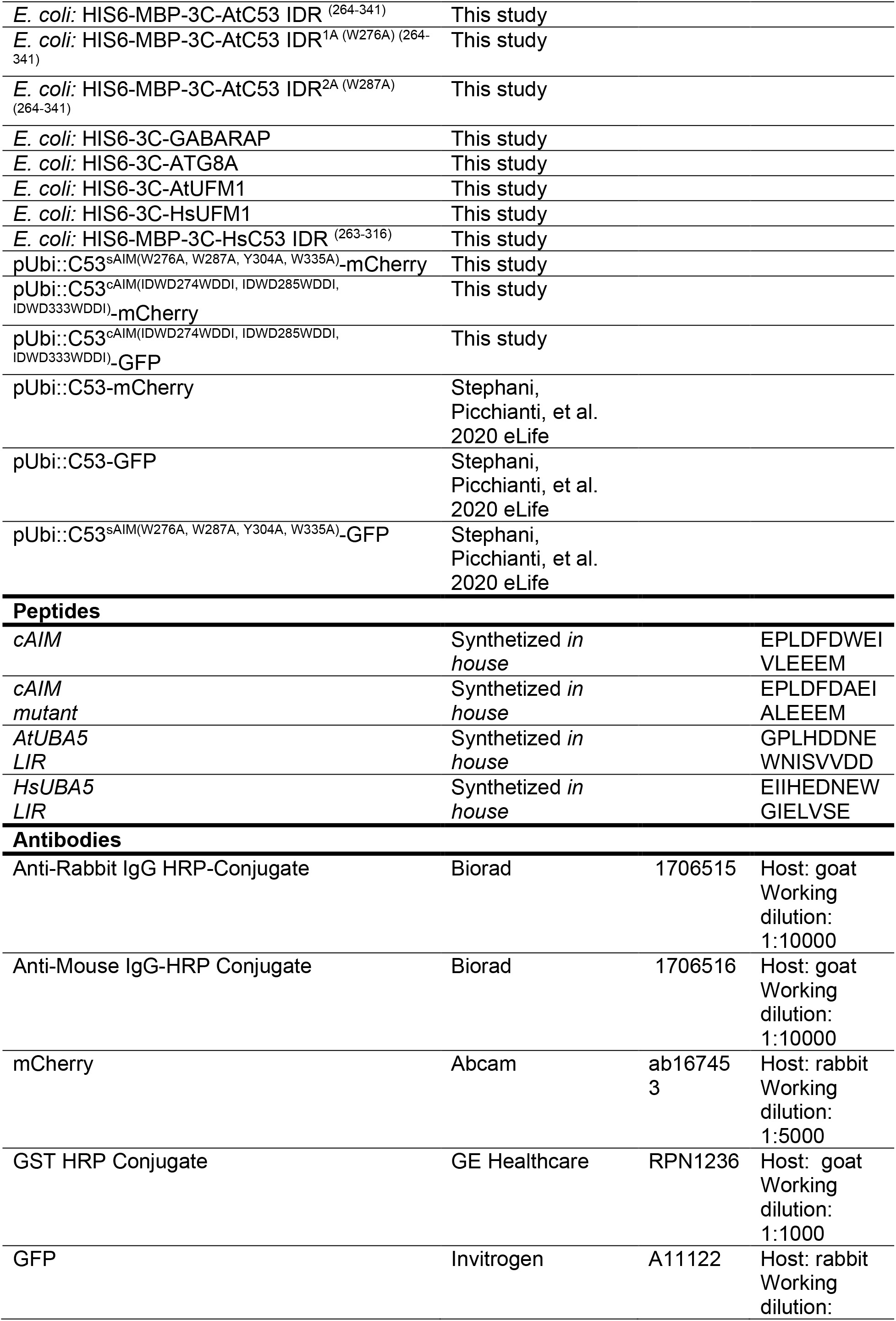
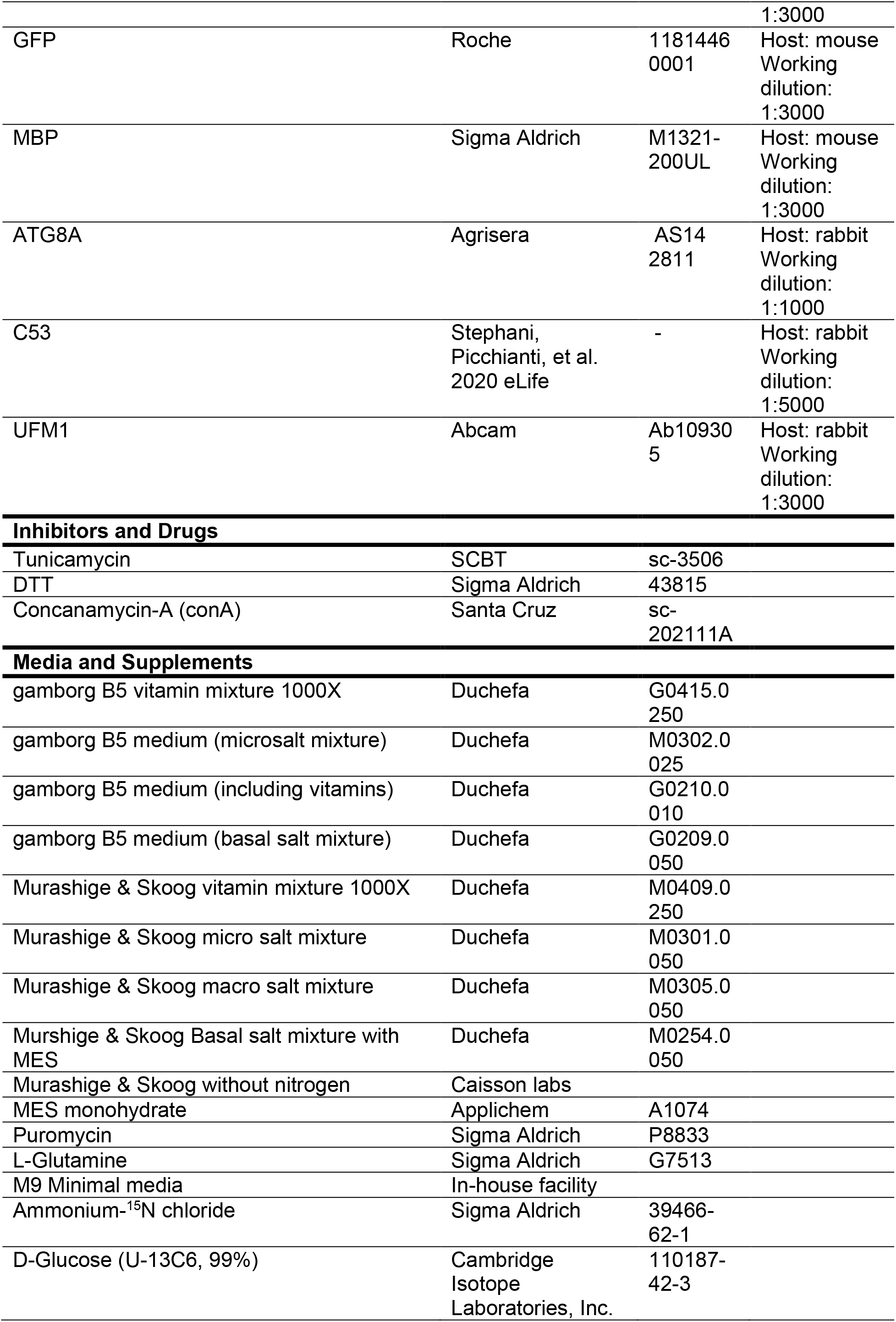
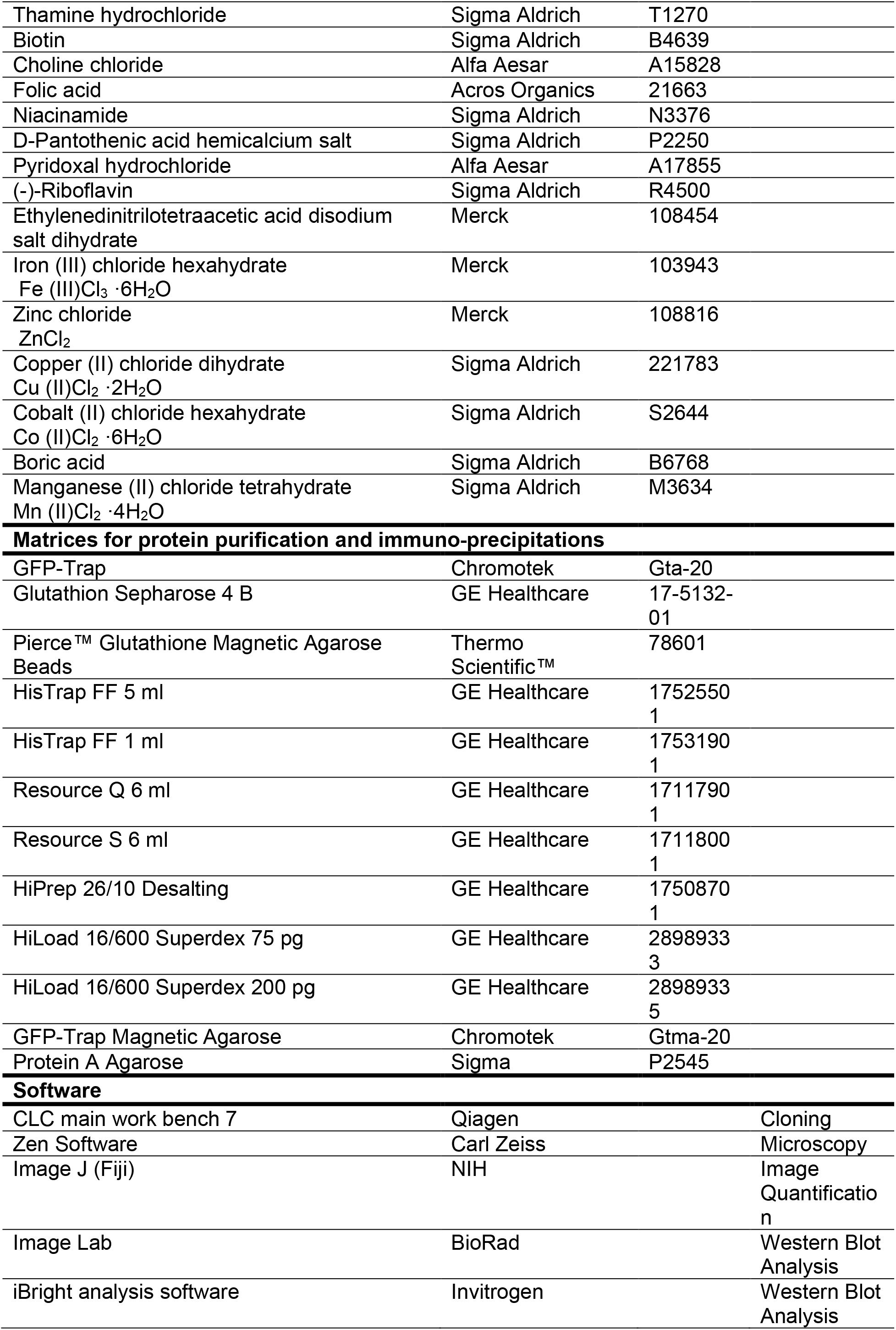
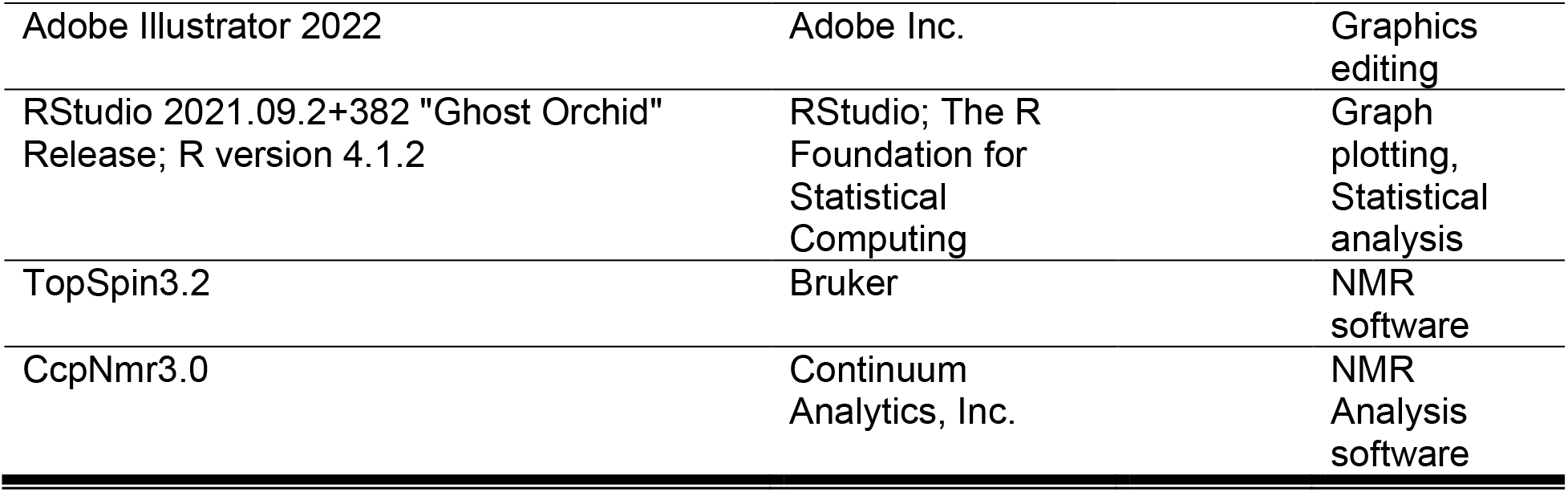

---

**Figure S1.**
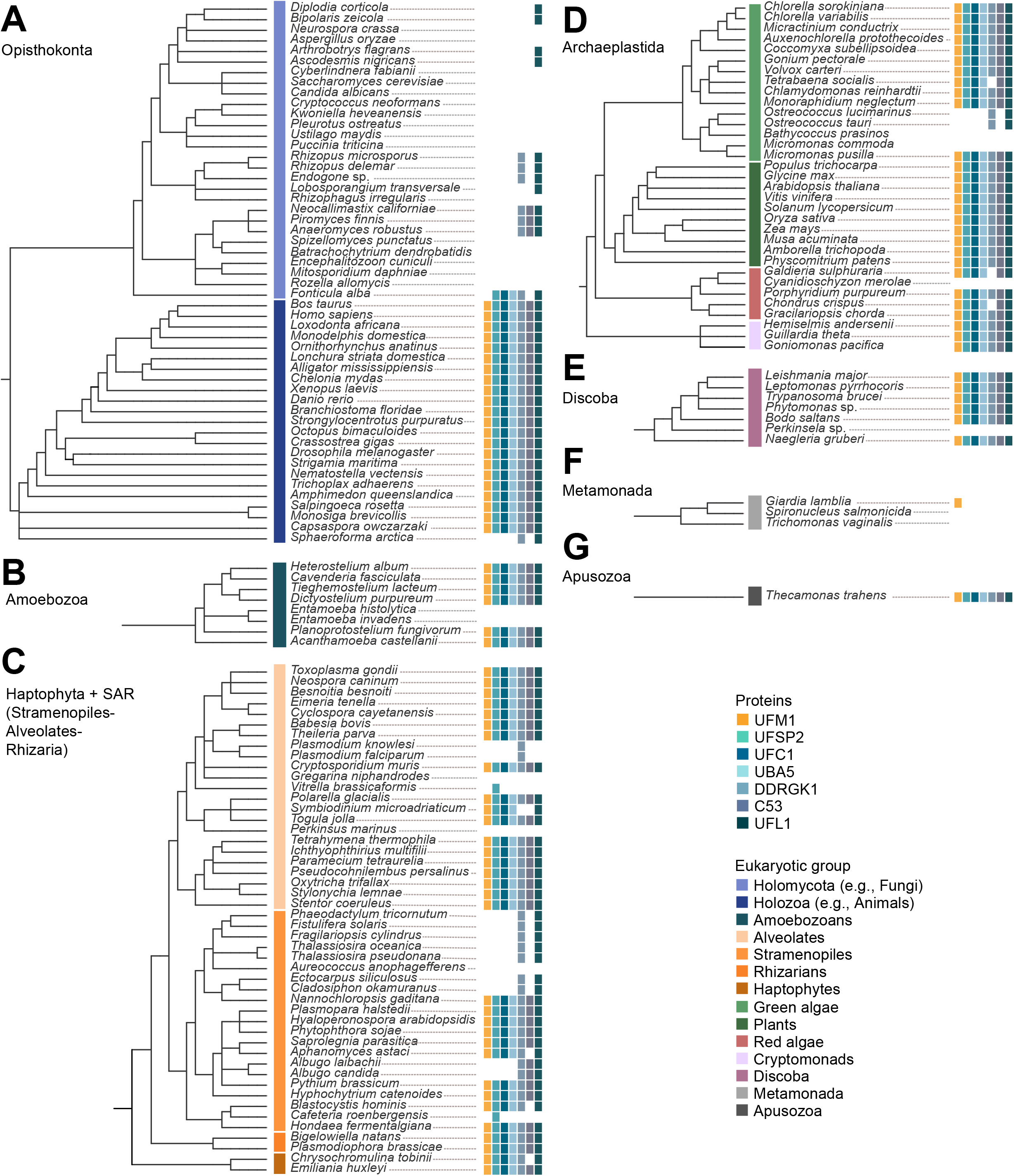
An expanded version of the tree depicted in Figure 1A, displaying the presence and absence of UFMylation proteins across the eukaryotic taxa. The tree has been divided into eukaryotic supergroups including the Opisthokonta (**A**), Amoebozoa (**B**), Haptophyta and SAR (**C**), Archaeplastida (**D**), Discoba (**E**), Metamonada (**F**), and Apusozoa (**G**).

**Figure S2.**
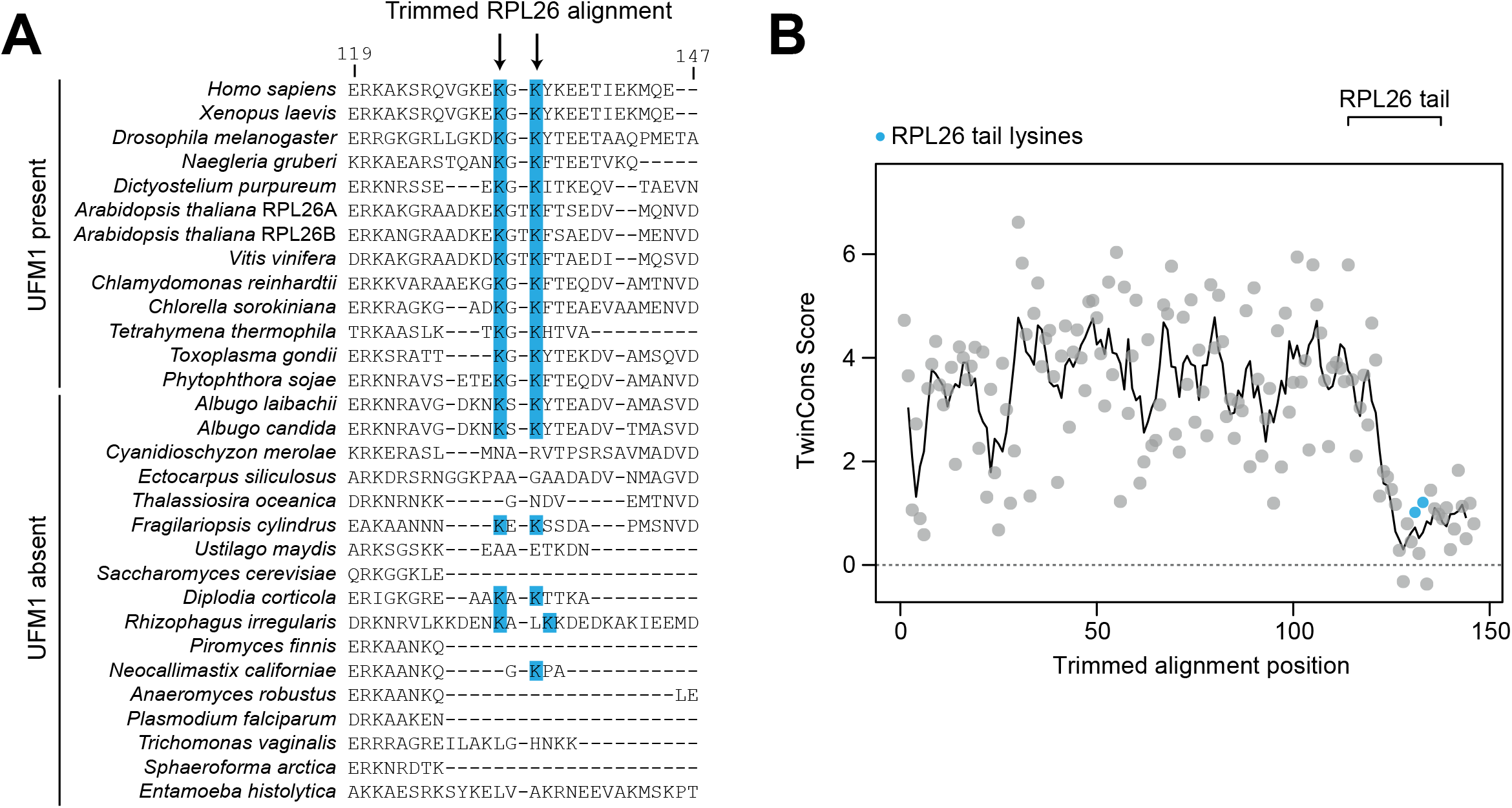
Conservation analysis of RPL26 shows that the ufmylated tail region is divergent. **(A) Multiple sequence alignment of RPL26 showing the conservation of the C-terminal tail in species with and without UFM1.** Lysine residues that are ufmylated have been highlighted. (**B) TwinCons analysis comparing the sequence conservation of RPL26.** The tail region is highly polymorphic.

**Figure S3.**
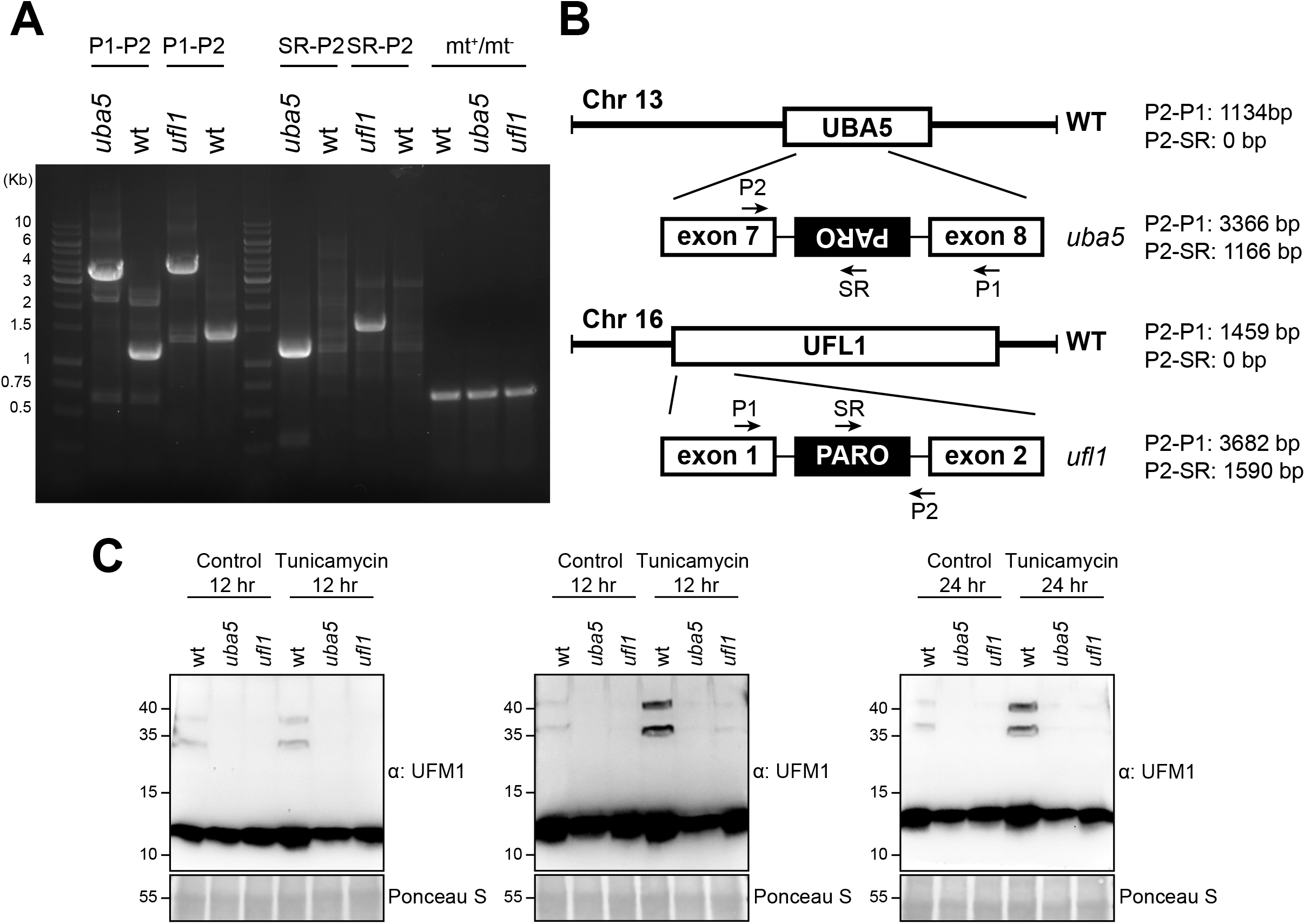
Characterization of the *Chlamydomonas reinhardtii* UFMylation pathway mutants. **(A) Genotyping of *C. reinhardtii uba5* and *ufl1* mutants**. *Left Panel*, mating type (mt +/-) and insertion site PCR products from purified genomic DNA samples prepared from wt, *uba5* and *ufl1* genotypes. PCR products were run on a 1% (w/v) agarose gel. DNA size markers are reported in Kb. **(B) Schematic diagram indicating the insertion site of the mutagenic cassette (PARO) in *ufl1* and *uba5* mutants.** Primers are indicated with arrows and expected PCR products from wild type and mutants are reported next to each respective diagram. **(C) RPL26 mono- and di-UFMylation is lost in *uba5* and *ufl1* mutants**. Cells were either left untreated or treated for 24 hours with 200 ng/mL tunicamycin. Protein extracts were analyzed by immunoblotting with anti-UFM1 antibodies. Total proteins were analyzed by Ponceau S staining. Quantification is shown in Figure 1C.

**Figure S4.**
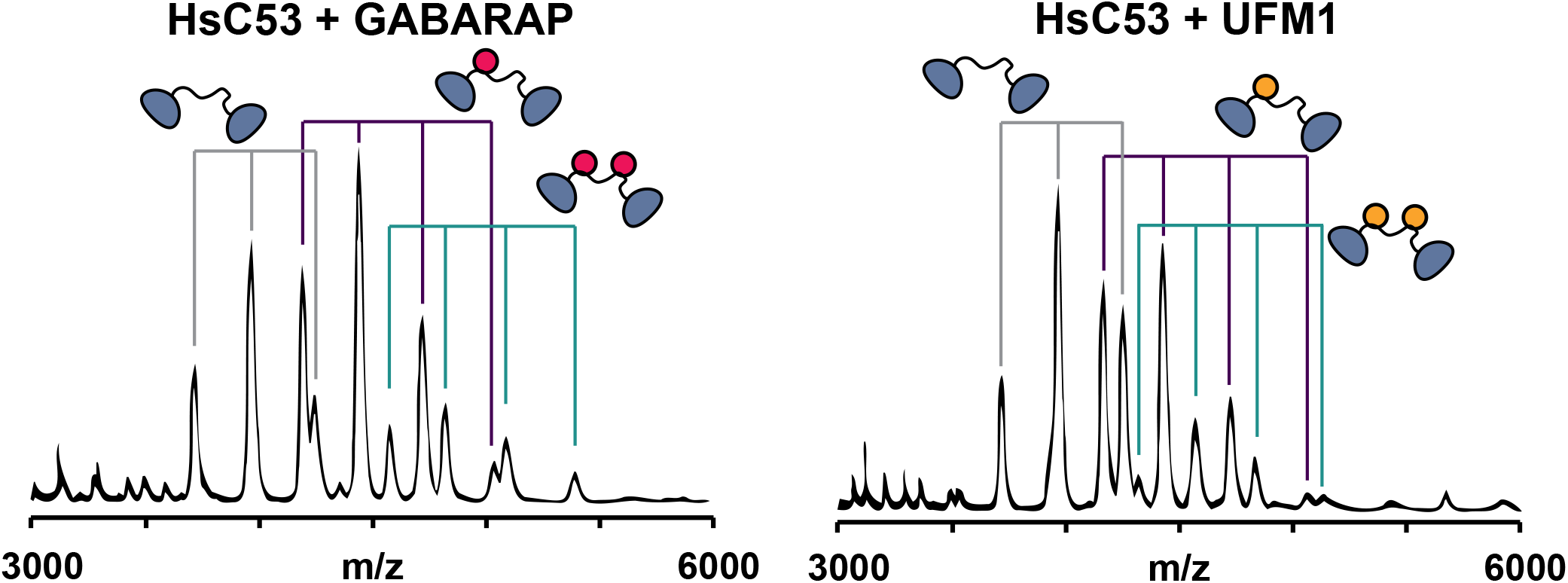
Native Mass-Spectrometry (nMS) spectra of HsC53 with GABARAP or HsUFM1 show very similar binding profiles. *Upper Panel*, GABARAP (4 µM) and HsC53 (2 µM). *Right Panel*, HsUFM1 (4 µM) and HsC53 (2 µM). Binding of HsC53 to GABARAP and HsUFM1 is observed in 1:1 (violet) and 1:2 ratios (teal).

**Figure S5.**
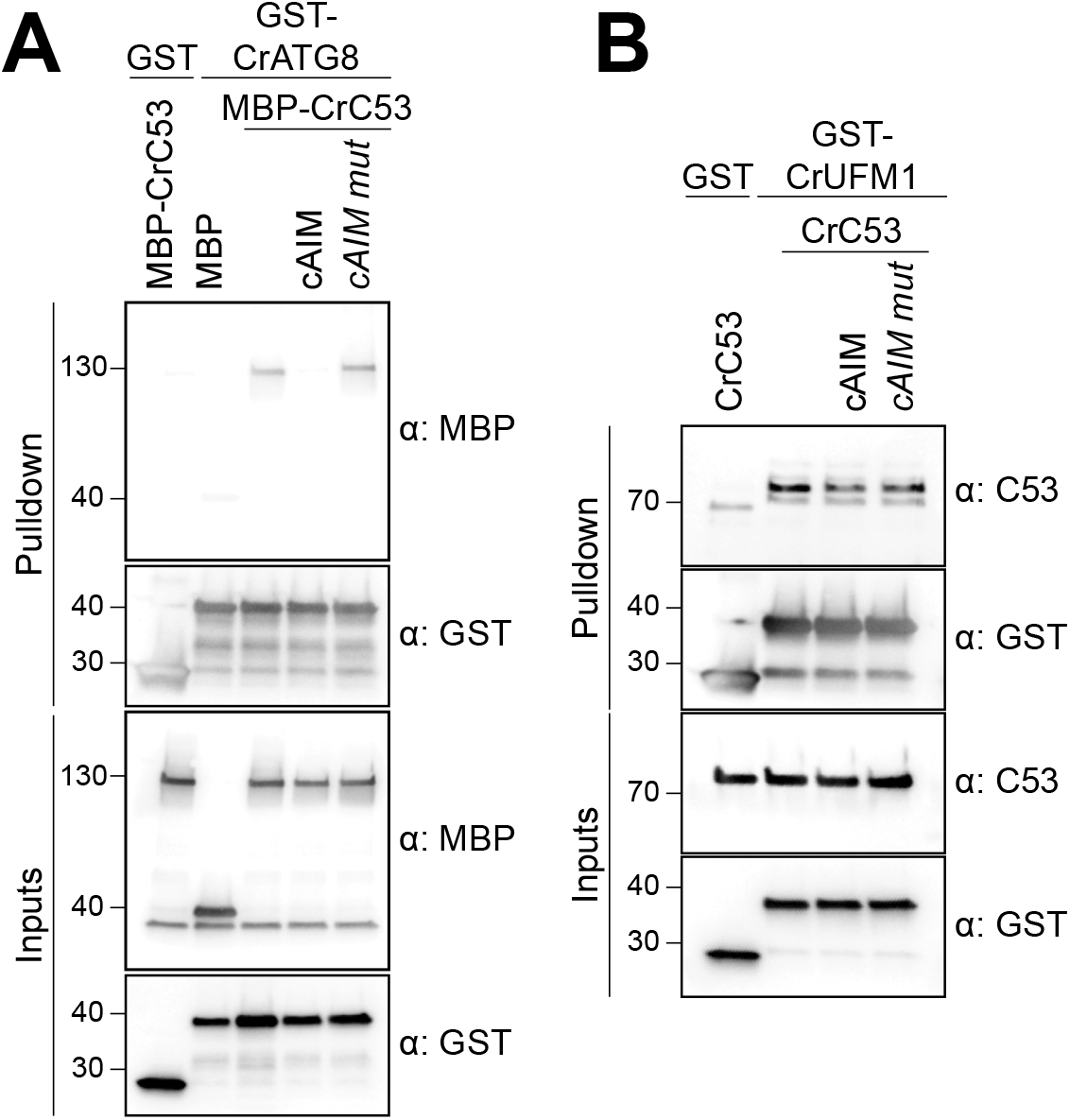
The canonical ATG8 Interacting Motif (cAIM) peptide cannot outcompete C53-UFM1 interaction for *C. reinhardtii* (Cr). **(A) CrC53 binds CrATG8A in a cAIM-dependent manner. (B) CrC53 binds UFM1 in a cAIM-independent manner.** Bacterial lysates containing recombinant protein or purified recombinant proteins were mixed and pulled down with glutathione magnetic agarose beads. Input and bound proteins were visualized by immunoblotting with anti-GST, anti-MBP or anti-AtC53 antibodies. cAIM wild type or mutant peptides were used to a final concentration of 200 µM.

**Figure S6.**
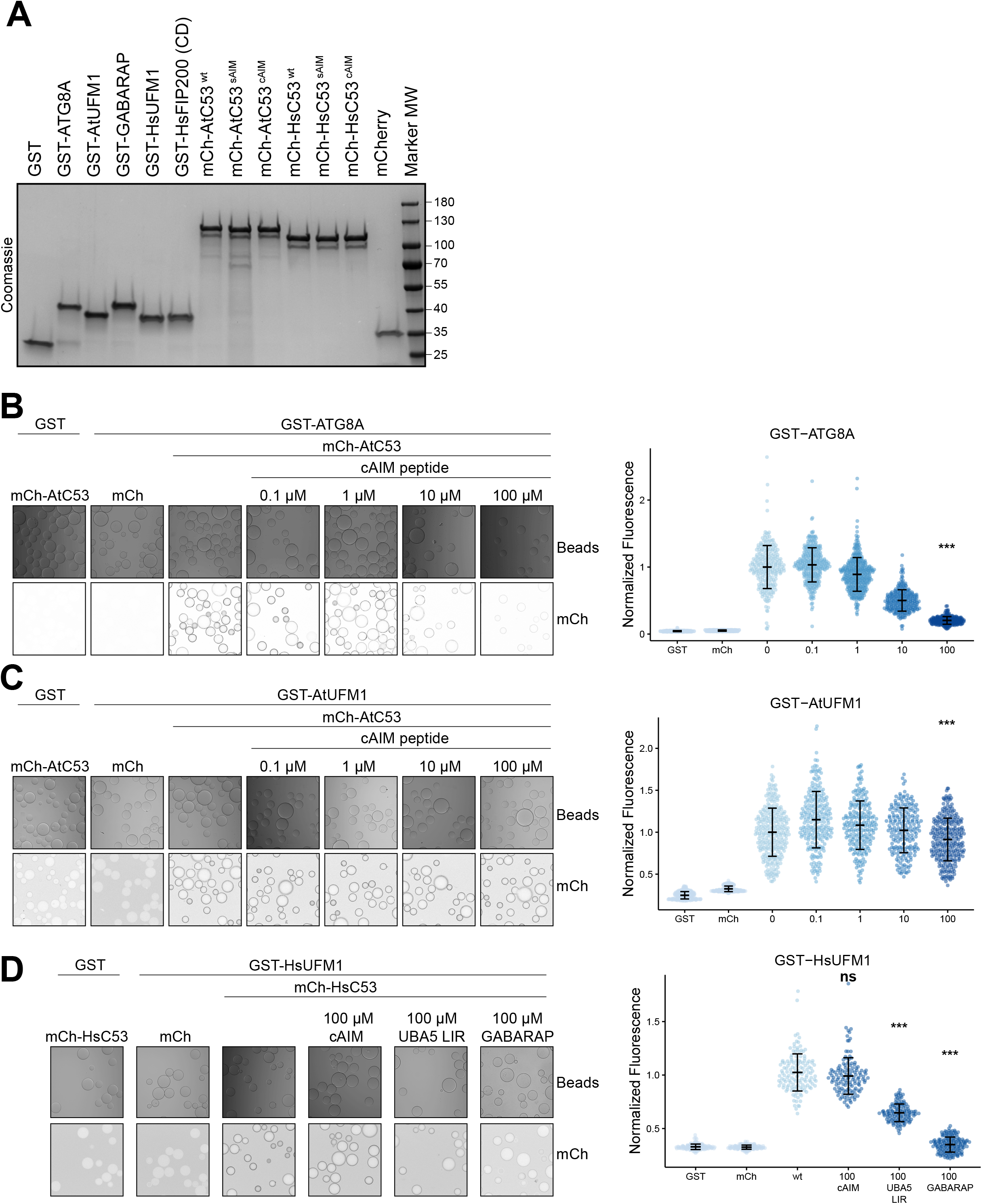
The canonical ATG8 Interacting Motif (cAIM) cannot outcompete C53-UFM1 interaction. **(A) Purified proteins used for the protein-protein interaction microscopy binding assays.** Recombinant proteins were analyzed for purity by SDS-PAGE followed by Coomassie staining. Marker molecular weights (MW) are indicated in kDa. mCh: mCherry. **(B, C) Microscopy-based protein–protein interaction assays showing unlike ATG8A-C53 interaction, UFM1-C53 interaction is insensitive to cAIM peptide competition**. Glutathione-sepharose beads were prepared by incubating them with GST-ATG8A (C) or GST-AtUFM1 (D). The pre-assembled beads were then washed and mixed with 1 µM of AtC53 containing increasing concentrations of cAIM peptide (0-100 µM). The beads were then imaged using a confocal microscope. *Left Panel,* representative confocal images (inverted grayscale) for each condition are shown. *Right panel*, normalized fluorescence is shown for each condition with the mean (± SD) of 2 independent replicates containing 2 technical replicates. Unpaired two-samples Wilcoxon test with continuity correction was performed to analyze the differences between wild type without cAIM peptide and wild type with 100 µM cAIM peptide. *, p-value < 0.05, ***, p-value < 0.001. Total number of beads, mean, median, standard deviation and p-values are reported in Supplementary data 2. **(D) Microscopy-based protein–protein interaction assays showing UBA5 LIR peptide and GABARAP can compete for C53 interaction with UFM1**. Glutathione-sepharose beads were prepared by incubating them with GST-HsUFM1. The pre-assembled beads were then washed and mixed with 1 µM of HsC53 with either 100 µM cAIM peptide, 100 µM UBA5 LIR peptide or 100 µM GABARAP. The beads were then imaged using a confocal microscope. *Left Panel,* representative confocal images (inverted grayscale) for each condition are shown. *Right panel*, normalized fluorescence is shown for each condition with the mean (± SD). Unpaired two-samples Wilcoxon test with continuity correction was performed to analyze the differences between wild type and wild type mixed with either cAIM peptide, UBA5 LIR peptide or GABARAP. ns, p-value > 0.05, ***, p-value < 0.001. Total number of beads, mean, median, standard deviation and p-values are reported in Supplementary data 2.

**Figure S7.**
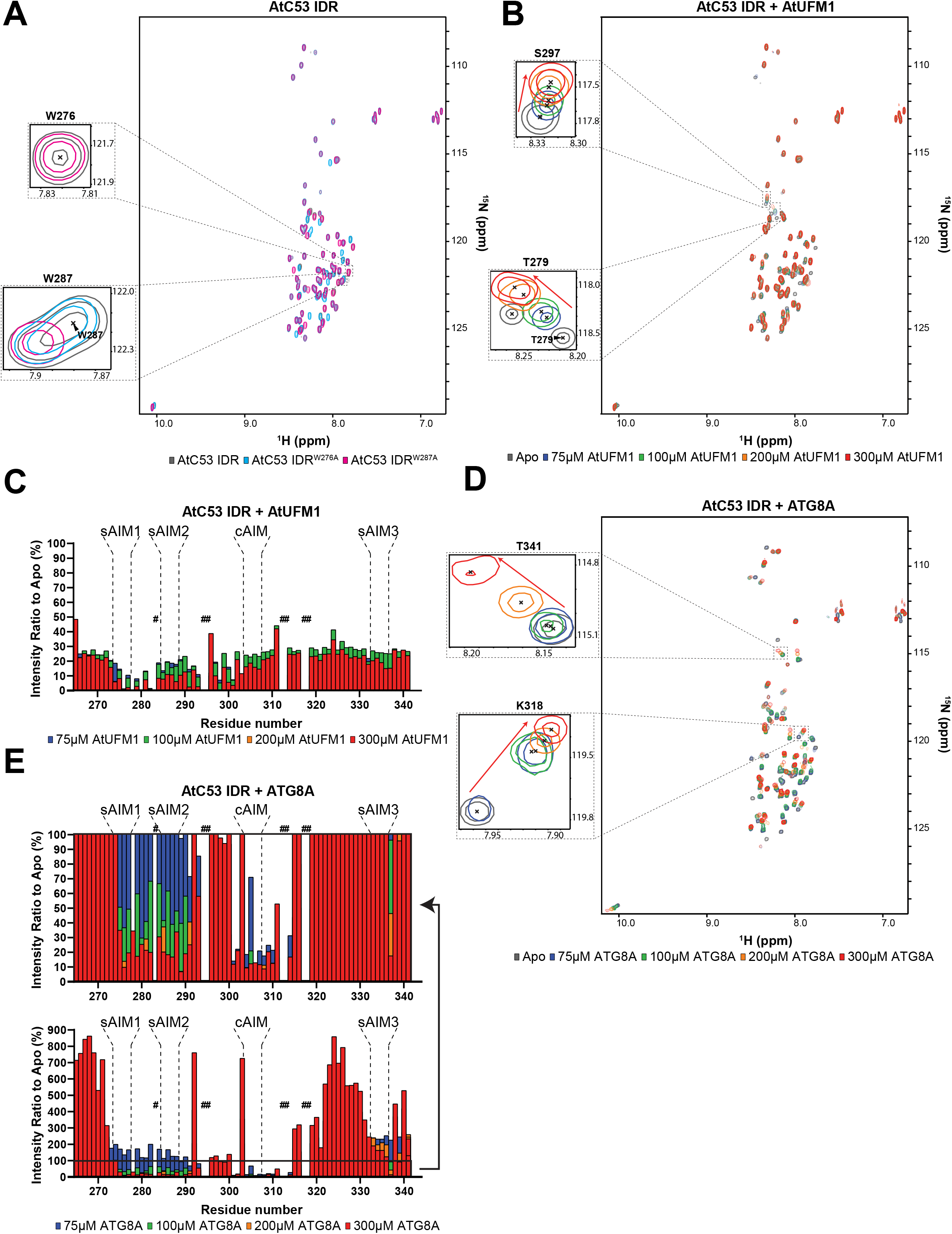
Structural characterization of AtC53 IDR binding to AtUFM1 and ATG8A using NMR spectroscopy. **(A) Validation of AtC53 IDR backbone resonance assignments.** Overlaid ^1^H-^15^N HSQC spectra of isotope-labeled AtC53 IDR (grey), AtC53 IDR^W276A^ (cyan) and AtC53 IDR^W287A^ (magenta). Insets of resonances corresponding to residues W276 and W287 are shown. **(B) Addition of AtUFM1 changes the magnetic resonance of specific residues in AtC53.** Overlaid ^1^H-^15^N HSQC spectra of isotope-labeled AtC53 IDR in their free (gray) or bound state to 75 µM (blue), 100 µM (green), 200 µM (orange) and 300 µM (red) unlabeled AtUFM1. Examples of individual peaks that shift upon binding are shown as insets. Chemical shifts are indicated with arrows. **(C) Signal intensity changes in AtC53 IDR upon binding of AtUFM1 are concentration dependent.** Intensity ratio broadening of AtC53 IDR (100 µM) in the presence of 75 µM (blue), 100 µM (green), 200 µM (orange) and 300 µM (red) AtUFM1. Bars corresponding to residues in AIMs are highlighted. Unassigned AtC53 IDR residues are indicated by hashtags. **(D) Addition of ATG8A affects a greater number of residues in the AtC53 IDR spectra.** Overlaid ^1^H-^15^N HSQC spectra of isotope-labeled AtC53 IDR in their free (gray) or bound state to 75 µM (blue), 100 µM (green), 200 µM (orange) and 300 µM (red) unlabeled ATG8A. Insets of individual peaks that shifted upon binding are shown. Chemical shifts are indicated with arrows. **(E) Signal intensity changes in AtC53 IDR upon binding of ATG8A are concentration dependent.** Intensity ratio broadening of AtC53 IDR (100 µM) in the presence of 75 µM (blue), 100 µM (green), 200 µM (orange) and 300 µM (red) ATG8A. Top panel represents an inset of lower panel. Unassigned AtC53 IDR residues are indicated by hashtags. Bars corresponding to residues in AIMs are highlighted.

**Figure S8.**
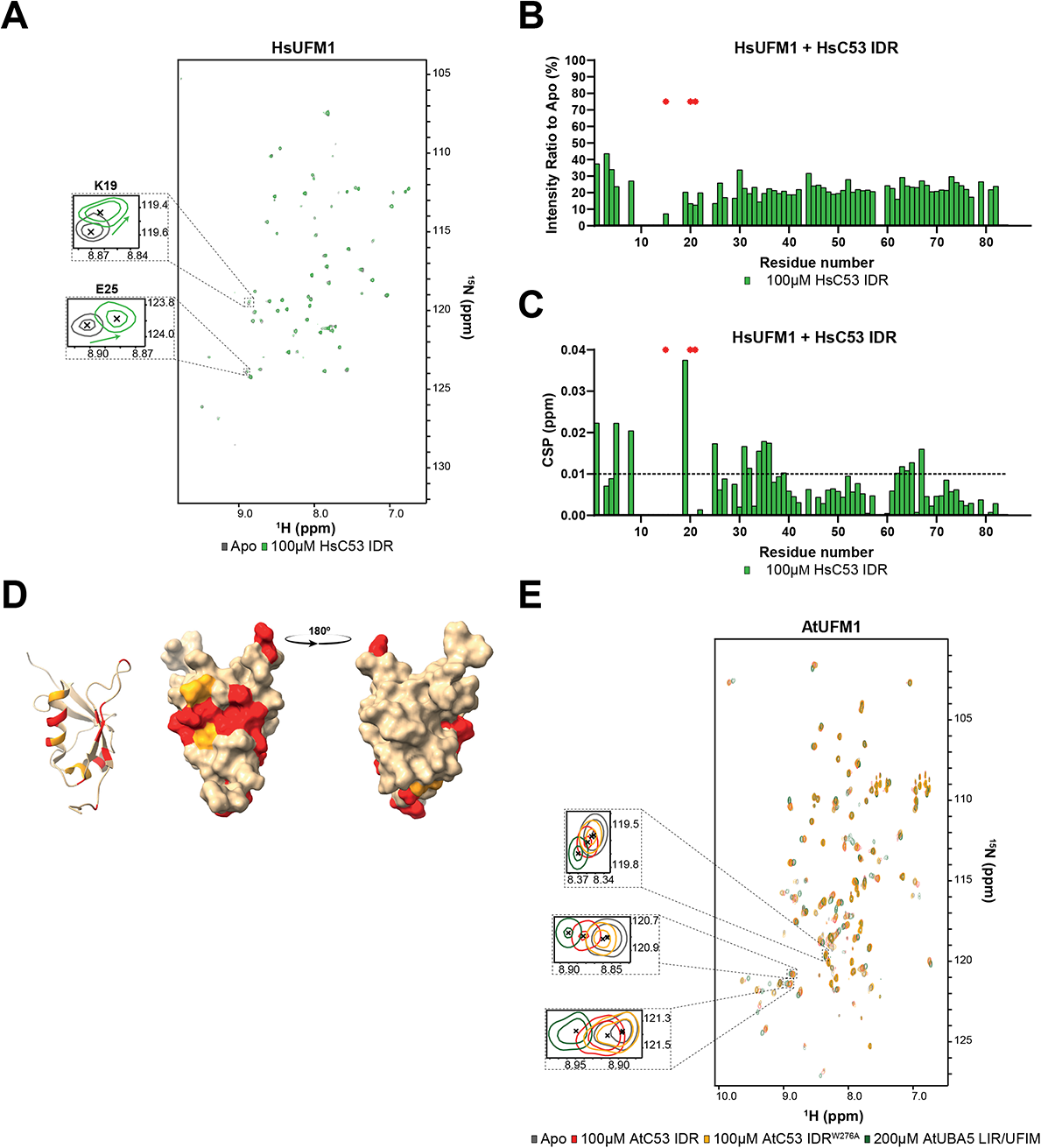
Structural characterization of UFM1 binding to C53 IDR using NMR spectroscopy. **(A) A small number of residues are affected by the addition of HsC53 IDR as shown in the HsUFM1 spectra.** Overlaid ^1^H-^15^N HSQC spectra of isotope-labeled HsUFM1 in their free (gray) or bound state to 100 µM unlabeled HsC53 IDR (green). Insets of individual peaks that shift upon binding are shown. **(B) HsC53 IDR binding to HsUFM1 causes general signal intensity drop in HsUFM1 spectra.** Intensity ratio broadening of HsUFM1 (100 µM) in the presence of 100 µM HsC53 IDR (green). HN resonances for residues that could not be assigned in the bound state are shown as red asterisks. **(C) Chemical shift perturbations (CSPs) in the HsUFM1 spectrum (grey) upon addition of 100 µM HsC53 IDR (green).** HN resonances for residues that could not be assigned in the bound state are shown as red asterisks. The dashed line represents S.D. **(D) Three-dimensional mapping of residues showing CSP in HsUFM1 NMR spectra upon HsC53 IDR binding.** CSPs were mapped on the UFM1 structure (PDB: 1WXS) presented schematically on the left plot and as a surface representation in two projections on the right plot. Residues that are not affected or are slightly (CSP < 0.01), intermediately (0.01 < CSP < 0.015), or strongly (CSP > 0.015) affected by the binding are colored in tan, orange and red, respectively. **(E) AtC53 IDR binding to AtUFM1 is similar to that of AtUBA5 and involves sAIM1.** Overlaid ^1^H-^15^N HSQC spectra of isotope-labeled AtUFM1 in their free (gray) or bound state to 100 µM unlabeled AtC53 IDR (red), 100 µM unlabeled AtC53 IDR^W276A^ (yellow) or AtUBA5 LIR/UFIM (green). Insets of chemical shift perturbations of individual peaks are shown.

**Figure S9.**
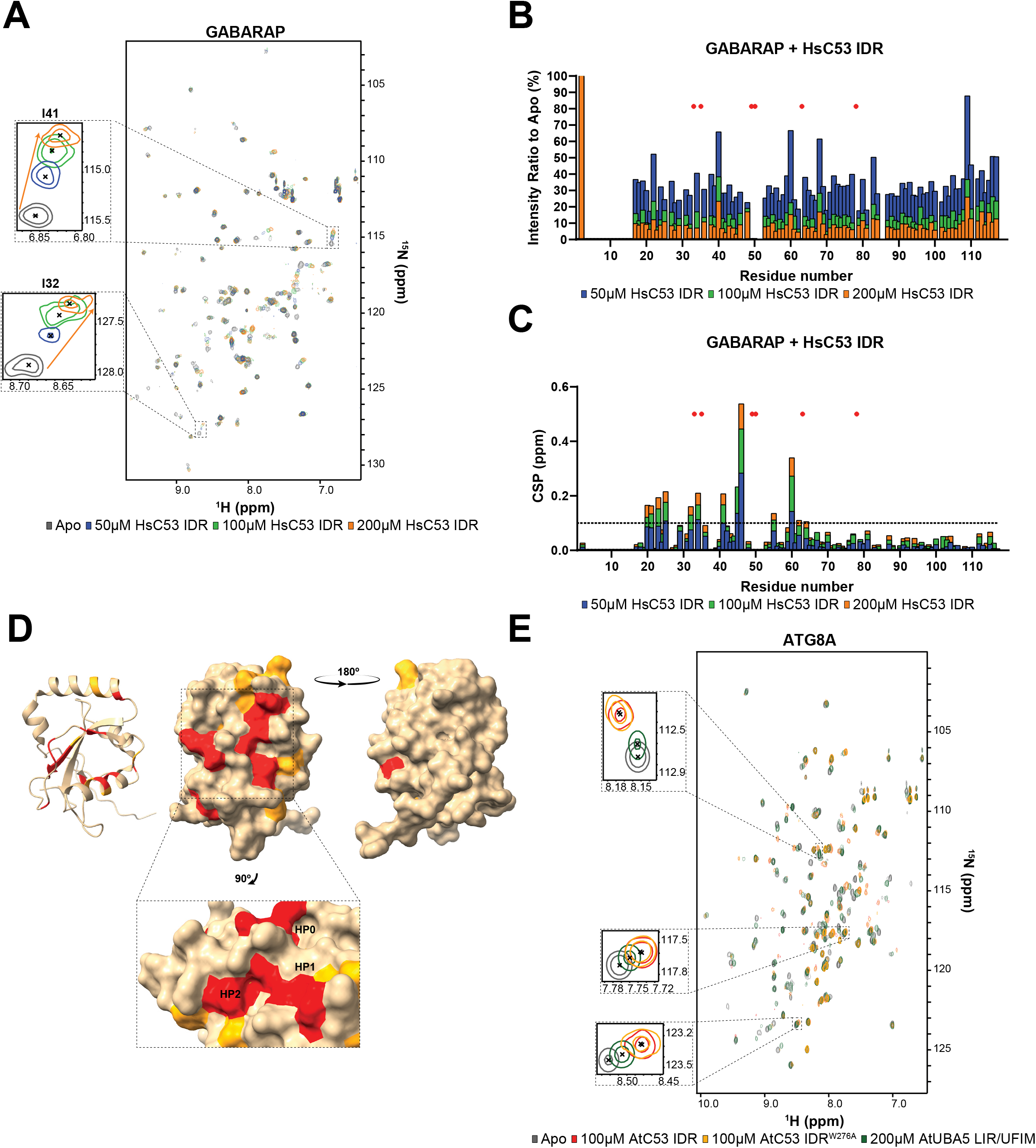
Structural characterization of ATG8 binding to C53 IDR using NMR spectroscopy. **(A) Addition of HsC53 IDR affects numerous residues in the GABARAP spectra.** Overlaid ^1^H-^15^N HSQC spectra of isotope-labeled GABARAP in their free (gray) or bound state to 50 µM (blue), 100 µM (green) or 200 µM (orange) unlabeled HsC53 IDR. Insets of individual peaks that shifted upon binding are shown. **(B) HsC53 IDR binding to GABARAP causes a general signal intensity drop in GABARAP spectra.** Intensity ratio broadening of GABARAP (100µM) in the presence of 50 µM (blue), 100 µM (green) or 200 µM (orange) unlabeled HsC53 IDR. HN resonances for residues that could not be assigned in the bound state are shown as red asterisks. **(C) NMR chemical shift perturbations (CSP) of GABARAP in the presence of 50 µM (blue), 100 µM (green) or 200 µM (orange) HsC53 IDR.** HN resonances for residues that could not be assigned in the bound state are shown as red asterisks. The dashed line represents S.D. **(D) Three-dimensional mapping of residues showing CSP in GABARAP NMR spectra upon HsC53 IDR binding.** CSPs were mapped on the GABARAP structure (PDB: 6HB9) presented schematically on the left plot and as a surface representation in two projections on the right plot. Residues that are not affected or are slightly (CSP < 0.1), intermediately (0.1 < CSP < 0.2), or strongly (CSP > 0.2) affected by the binding are colored in tan, orange and red, respectively. The inset highlights the position of the HP0, HP1 and HP2 hydrophobic pockets in GABARAP. **(E) AtC53 IDR binding to ATG8 is similar to that of AtUBA5.** Overlaid ^1^H-^15^N HSQC spectra of isotope-labeled ATG8A in their free (gray) or bound state to 100 µM unlabeled AtC53 IDR (red), 100 µM unlabeled AtC53 IDR^W276A^ (yellow) or 200 µM AtUBA5 LIR/UFIM (green). Insets of chemical shift perturbations of individual peaks are shown.

**Figure S10.**
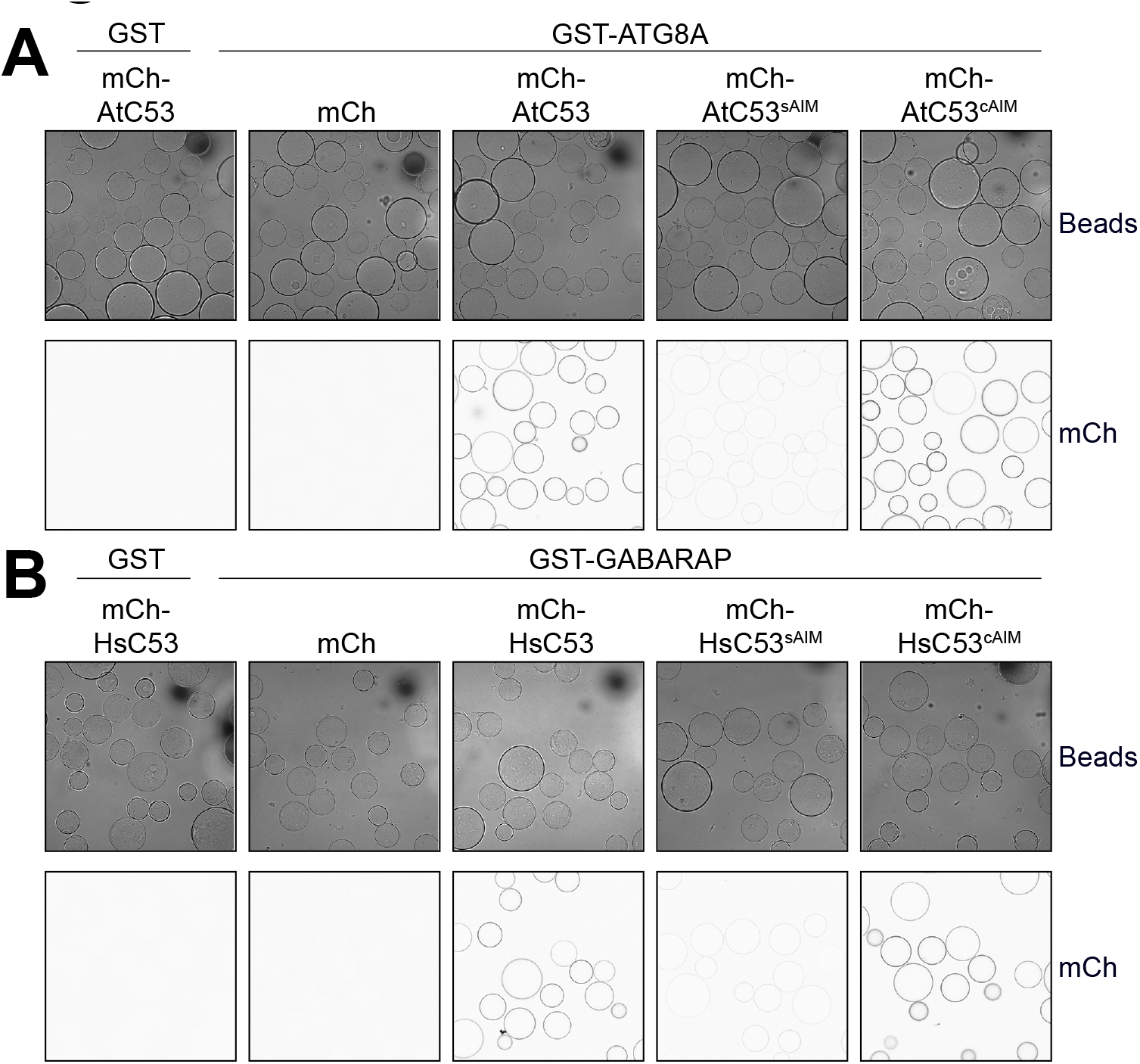
Microscopy-based protein–protein interaction assays showing C53^cAIM^ has increased affinity towards ATG8 or GABARAP. **(A, B)** Representative confocal images (inverted grayscale) for each condition from Figure 5 D, E are shown.

**Fig. S11.**
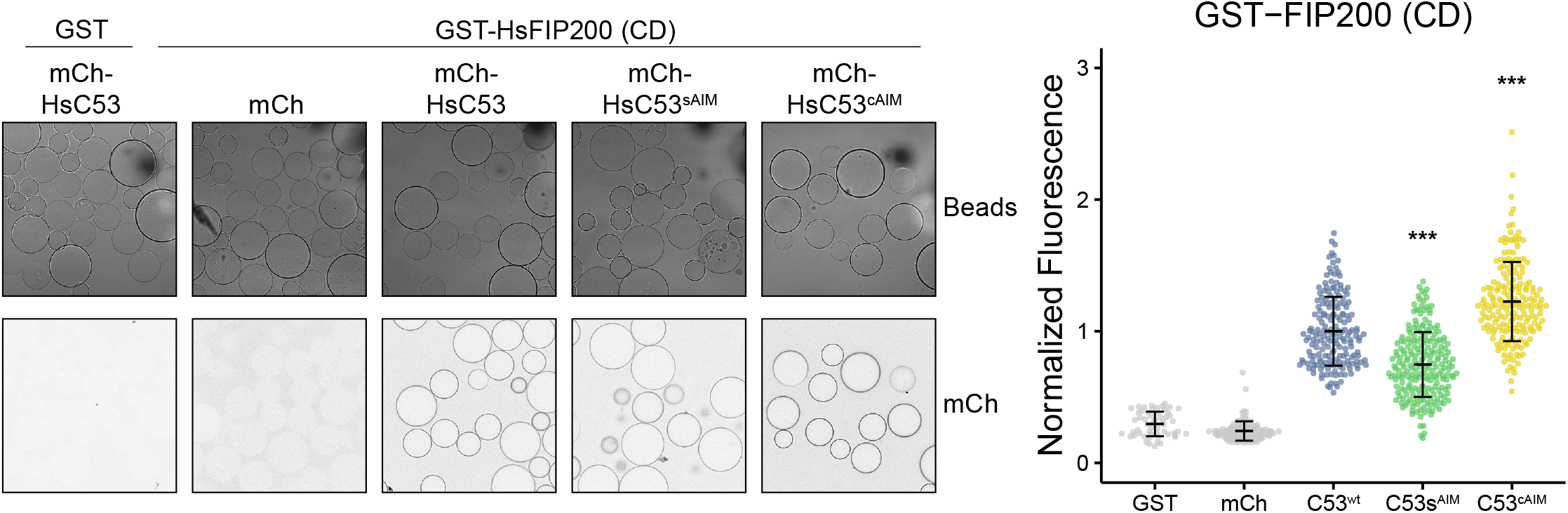
C53-HsFIP200 Claw domain (CD) interaction is also mediated by the sAIM sequences and strengthened by sAIM to cAIM conversion. Glutathione-sepharose beads were prepared by incubating them with GST-FIP200 CD. The pre-assembled beads were then washed and mixed with 1 µM of HsC53, 1 µM of HsC53^sAIM^ or 1 µM of HsC53^cAIM^ mutants. The beads were then imaged using a confocal microscope. *Left Panel,* representative confocal images (inverted grayscale) for each condition are shown. *Right panel*, normalized fluorescence is shown for each condition with the mean (± SD) of 2 independent replicates containing 2 technical replicates. Unpaired two-samples Wilcoxon test with continuity correction was performed to analyze the differences between wild type and mutants. ***, p-value < 0.001. Total number of beads, mean, median, standard deviation and p-values are reported in Supplementary data 2.

**Figure S12.**
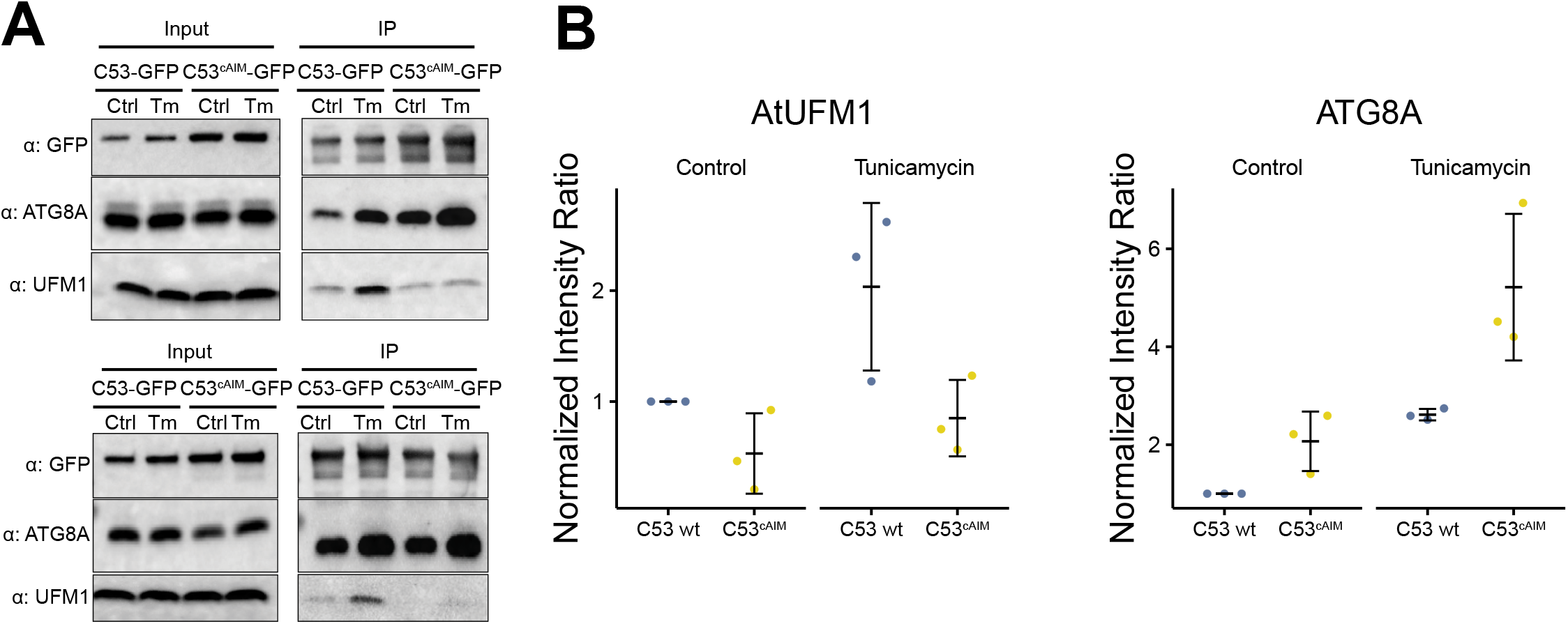
*In vivo* pull downs showing sAIM to cAIM conversion strengthening C53-ATG8 association and weakening C53-UFM1 association. **(A) Biological replicates of representative experiment shown in Figure5F.** 6-day old *Arabidopsis* seedlings expressing AtC53-GFP, AtC53^cAIM^-GFP in *c53* mutant background were incubated in liquid 1/2 MS medium with 1% sucrose supplemented with DMSO as control (Ctrl) or 10 μg/ml tunicamycin (Tm) for 16 hours and used for co-immunoprecipitation. Lysates were incubated with GFP-Trap Magnetic Agarose, input and bound proteins were detected by immunoblotting using the respective antibodies as indicated. **(B)** Quantification of blots in (Fig. 5F, Fig. S12A), UFM1 and ATG8 protein levels that associate with AtC53-GFP or AtC53^cAIM^-GFP are shown. Bars represent the mean (± SD) of 3 biological replicates (BR).

**Supplementary Data S1.** Eukaryotic datasets used in the phylogenomic analysis. Species names, NCBI Taxonomy identifiers, genome assemblies, proteomes, and their sources for each species analyzed are provided.

**Supplementary Data S2.** Total number of beads, mean, median, standard deviation and p-values of the microscopy-based protein-protein interaction assays are reported.

**Supplementary Data S3.** Fiji macro and agarose bead model for automatic quantification.

## References

1. Pohl C, Dikic I (2019) Cellular quality control by the ubiquitin-proteasome system and autophagy. Science 366:818–822. https://doi.org/10.1126/science.aax3769

2. Sun Z, Brodsky JL (2019) Protein quality control in the secretory pathway. J Cell Biol 218:3171–3187. https://doi.org/10.1083/jcb.201906047

3. Karagöz GE, Acosta-Alvear D, Walter P (2019) The Unfolded Protein Response: Detecting and Responding to Fluctuations in the Protein-Folding Capacity of the Endoplasmic Reticulum. Cold Spring Harb Perspect Biol 11:. https://doi.org/10.1101/cshperspect.a033886

4. Kirkin V, Rogov VV (2019) A Diversity of Selective Autophagy Receptors Determines the Specificity of the Autophagy Pathway. Molecular Cell 76:268–285. https://doi.org/10.1016/j.molcel.2019.09.005

5. Dikic I (2017) Proteasomal and Autophagic Degradation Systems. Annu Rev Biochem 86:193–224. https://doi.org/10.1146/annurev-biochem-061516-044908

6. Klionsky DJ, Petroni G, Amaravadi RK, et al (2021) Autophagy in major human diseases. EMBO J 40:e108863. https://doi.org/10.15252/embj.2021108863

7. Stephani M, Dagdas Y (2020) Plant Selective Autophagy—Still an Uncharted Territory With a Lot of Hidden Gems. J Mol Biol 432:. https://doi.org/10.1016/j.jmb.2019.06.028

8. Marshall RS, Vierstra RD (2018) Autophagy: The Master of Bulk and Selective Recycling. Annu Rev Plant Biol 69:173–208. https://doi.org/10.1146/annurev-arplant-042817-040606

9. Mizushima N (2018) A brief history of autophagy from cell biology to physiology and disease. Nat Cell Biol 20:521–527. https://doi.org/10.1038/s41556-018-0092-5

10. Dikic I, Elazar Z (2018) Mechanism and medical implications of mammalian autophagy. Nat Rev Mol Cell Biol 19:349–364. https://doi.org/10.1038/s41580-018-0003-4

11. Kirkin V (2020) History of the Selective Autophagy Research: How Did It Begin and Where Does It Stand Today? Journal of Molecular Biology 432:3–27. https://doi.org/10.1016/j.jmb.2019.05.010

12. Birgisdottir ÅB, Lamark T, Johansen T (2013) The LIR motif - crucial for selective autophagy. Journal of Cell Science 126:3237–3247. https://doi.org/10.1242/jcs.126128

13. Johansen T, Lamark T (2020) Selective Autophagy: ATG8 Family Proteins, LIR Motifs and Cargo Receptors. Journal of Molecular Biology 432:80–103. https://doi.org/10.1016/j.jmb.2019.07.016

14. Stephani M, Picchianti L, Gajic A, et al (2020) A cross-kingdom conserved er-phagy receptor maintains endoplasmic reticulum homeostasis during stress. eLife 9:1–105. https://doi.org/10.7554/ELIFE.58396

15. Liang JR, Lingeman E, Luong T, et al (2020) A Genome-wide ER-phagy Screen Highlights Key Roles of Mitochondrial Metabolism and ER-Resident UFMylation. Cell 180:1160–1177.e20. https://doi.org/10.1016/j.cell.2020.02.017

16. Daniel J, Liebau E (2014) The Ufm1 Cascade. Cells 3:627–638. https://doi.org/10.3390/cells3020627

17. Banerjee S, Kumar M, Wiener R (2020) Decrypting UFMylation: How Proteins Are Modified with UFM1. Biomolecules 10:. https://doi.org/10.3390/biom10101442

18. Oweis W, Padala P, Hassouna F, et al (2016) Trans-Binding Mechanism of Ubiquitin-like Protein Activation Revealed by a UBA5-UFM1 Complex. Cell Reports 16:3113– 3120. https://doi.org/10.1016/j.celrep.2016.08.067

19. Kumar M, Padala P, Fahoum J, et al (2021) Structural basis for UFM1 transfer from UBA5 to UFC1. Nature Communications 12:1–13. https://doi.org/10.1038/s41467-021-25994-6

20. Zhou J, Liang Q, Dong M, et al (2022) Optimized protocol to detect protein UFMylation in cells and in vitro via immunoblotting. STAR Protocols 3:101074. https://doi.org/10.1016/j.xpro.2021.101074

21. Peter JJ, Magnussen HM, DaRosa PA, et al (2022) Non-canonical scaffold-type ligase complex mediates protein UFMylation. bioRxiv 2022.01.31.478489

22. Walczak CP, Leto DE, Zhang L, et al (2019) Ribosomal protein RPL26 is the principal target of UFMylation. Proc Natl Acad Sci U S A 116:1299–1308. https://doi.org/10.1073/pnas.1816202116

23. Wang L, Xu Y, Rogers H, et al (2020) UFMylation of RPL26 links translocation-associated quality control to endoplasmic reticulum protein homeostasis. Cell Res 30:5– 20. https://doi.org/10.1038/s41422-019-0236-6

24. Wolf YI, Koonin EV (2013) Genome reduction as the dominant mode of evolution. BioEssays 35:829–837. https://doi.org/10.1002/bies.201300037

25. Ashkenazy H, Abadi S, Martz E, et al (2016) ConSurf 2016: an improved methodology to estimate and visualize evolutionary conservation in macromolecules. Nucleic Acids Res 44:W344–50. https://doi.org/10.1093/nar/gkw408

26. Habisov S, Huber J, Ichimura Y, et al (2016) Structural and Functional Analysis of a Novel Interaction Motif within UFM1-activating Enzyme 5 (UBA5) Required for Binding to Ubiquitin-like Proteins and Ufmylation. J Biol Chem 291:9025–9041. https://doi.org/10.1074/jbc.M116.715474

27. Huber J, Obata M, Gruber J, et al (2020) An atypical LIR motif within UBA5 (ubiquitin like modifier activating enzyme 5) interacts with GABARAP proteins and mediates membrane localization of UBA5. Autophagy 16:256–270. https://doi.org/10.1080/15548627.2019.1606637

28. Padala P, Oweis W, Mashahreh B, et al (2017) Novel insights into the interaction of UBA5 with UFM1 via a UFM1-interacting sequence. Scientific Reports 7:1–12. https://doi.org/10.1038/s41598-017-00610-0

29. Turco E, Witt M, Abert C, et al (2019) FIP200 Claw Domain Binding to p62 Promotes Autophagosome Formation at Ubiquitin Condensates. Mol Cell 74:330–346.e11. https://doi.org/10.1016/j.molcel.2019.01.035

30. Cox M, Dekker N, Boelens R, et al (1993) NMR studies of the POU-specific DNA-binding domain of Oct-1: Sequential proton and nitrogen-15 assignments and secondary structure. Biochemistry 32:6032–6040. https://doi.org/10.1021/bi00074a014

31. Sette M, Spurio R, van Tilborg P, et al (1999) Identification of the ribosome binding sites of translation initiation factor IF3 by multidimensional heteronuclear NMR spectroscopy. RNA 5:82–92. https://doi.org/10.1017/s1355838299981487

32. Wüthrich K (2003) NMR studies of structure and function of biological macromolecules (Nobel Lecture). J Biomol NMR 27:13–39. https://doi.org/10.1023/a:1024733922459

33. Zuiderweg ERP (2002) Mapping protein-protein interactions in solution by NMR spectroscopy. Biochemistry 41:1–7. https://doi.org/10.1021/bi011870b

34. Sasakawa H, Sakata E, Yamaguchi Y, et al (2006) Solution structure and dynamics of Ufm1, a ubiquitin-fold modifier 1. Biochem Biophys Res Commun 343:21–26. https://doi.org/10.1016/j.bbrc.2006.02.107

35. Kouno T, Miura K, Kanematsu T, et al (2002) Letter to the Editor: 1H, 13C and 15N resonance assignments of GABARAP, GABAA receptor associated protein. Journal of Biomolecular NMR 22:97–98. https://doi.org/10.1023/A:1013884402033

36. Noda NN, Ohsumi Y, Inagaki F (2010) Atg8-family interacting motif crucial for selective autophagy. FEBS Letters 584:1379–1385. https://doi.org/10.1016/j.febslet.2010.01.018

37. Zientara-Rytter K, Subramani S (2020) Mechanistic Insights into the Role of Atg11 in Selective Autophagy. J Mol Biol 432:104–122. https://doi.org/10.1016/j.jmb.2019.06.017

38. Turco E, Fracchiolla D, Martens S (2020) Recruitment and Activation of the ULK1/Atg1 Kinase Complex in Selective Autophagy. J Mol Biol 432:123–134. https://doi.org/10.1016/j.jmb.2019.07.027

39. Brandizzi F (2021) Maintaining the structural and functional homeostasis of the plant endoplasmic reticulum. Dev Cell. https://doi.org/10.1016/j.devcel.2021.02.008

40. Gerakis Y, Quintero M, Li H, Hetz C (2019) The UFMylation System in Proteostasis and Beyond. Trends Cell Biol 29:974–986. https://doi.org/10.1016/j.tcb.2019.09.005

41. Eck F, Kaulich M, Behrends C ACSL3 is a novel GABARAPL2 interactor that links ufmylation and lipid droplet biogenesis

42. Liu J, Guan D, Dong M, et al (2020) UFMylation maintains tumour suppressor p53 stability by antagonizing its ubiquitination. Nat Cell Biol 22:1056–1063. https://doi.org/10.1038/s41556-020-0559-z

43. Lee L, Perez Oliva AB, Martinez-Balsalobre E, et al (2021) UFMylation of MRE11 is essential for telomere length maintenance and hematopoietic stem cell survival. Sci Adv 7:eabc7371. https://doi.org/10.1126/sciadv.abc7371

44. Qin B, Yu J, Nowsheen S, et al (2019) UFL1 promotes histone H4 ufmylation and ATM activation. Nat Commun 10:1242. https://doi.org/10.1038/s41467-019-09175-0

45. Kulsuptrakul J, Wang R, Meyers NL, et al (2020) A genome-wide CRISPR screen identifies UFMylation and TRAMP-like complexes as host factors required for hepatitis A virus infection. Cold Spring Harbor Laboratory 2019.12.15.877100

46. UFMylation inhibits the proinflammatory capacity of interferon-γ–activated macrophages. In: PNAS. https://www.pnas.org/content/pnas/118/1/e2011763118. Accessed 4 Apr 2022

47. Tsaban T, Stupp D, Sherill-Rofe D, et al (2021) CladeOScope: Functional interactions through the prism of clade-wise co-evolution. NAR Genomics and Bioinformatics 3:1–13. https://doi.org/10.1093/nargab/lqab024

48. Buchfink B, Xie C, Huson DH (2014) Fast and sensitive protein alignment using DIAMOND. Nature Methods 12:59–60. https://doi.org/10.1038/nmeth.3176

49. Katoh K, Standley DM (2013) MAFFT multiple sequence alignment software version 7: Improvements in performance and usability. Molecular Biology and Evolution 30:772– 780. https://doi.org/10.1093/molbev/mst010

50. Capella-Gutiérrez S, Silla-Martínez JM, Gabaldón T (2009) trimAl: A tool for automated alignment trimming in large-scale phylogenetic analyses. Bioinformatics 25:1972–1973. https://doi.org/10.1093/bioinformatics/btp348

51. Minh BQ, Schmidt HA, Chernomor O, et al (2020) IQ-TREE 2: New models and efficient methods for phylogenetic inference in the genomic era. Molecular Biology and Evolution 37:1530–1534. https://doi.org/10.1093/molbev/msaa015

52. Boeckmann B, Bairoch A, Apweiler R, et al (2003) The SWISS-PROT protein knowledgebase and its supplement TrEMBL in 2003. Nucleic Acids Research 31:365–370. https://doi.org/10.1093/nar/gkg095

53. El-Gebali S, Mistry J, Bateman A, et al (2019) The Pfam protein families database in 2019. Nucleic Acids Research 47:D427–D432. https://doi.org/10.1093/nar/gky995

54. Rambaut A (2012) FigTree, a graphical viewer of phylogenetic trees. See http://tree.bio.ed.ac.uk/software/figtree

55. Rozewicki J, Li S, Amada KM, et al (2019) MAFFT-DASH: Integrated protein sequence and structural alignment. Nucleic Acids Research 47:W5–W10. https://doi.org/10.1093/nar/gkz342

56. Mistry J, Finn RD, Eddy SR, et al (2013) Challenges in homology search: HMMER3 and convergent evolution of coiled-coil regions. Nucleic Acids Research 41:e121. https://doi.org/10.1093/nar/gkt263

57. Slater GSC, Birney E (2005) Automated generation of heuristics for biological sequence comparison. BMC Bioinformatics 6:31. https://doi.org/10.1186/1471-2105-6-31

58. Letunic I, Bork P (2021) Interactive tree of life (iTOL) v5: An online tool for phylogenetic tree display and annotation. Nucleic Acids Research 49:W293–W296. https://doi.org/10.1093/nar/gkab301

59. Burki F, Roger AJ, Brown MW, Simpson AGB (2020) The new tree of eukaryotes. Trends in Ecology and Evolution 35:43–55. https://doi.org/10.1016/j.tree.2019.08.008

60. Federhen S (2012) The NCBI Taxonomy database. Nucleic Acids Research 40:136–143. https://doi.org/10.1093/nar/gkr1178

61. Edgar RC (2004) MUSCLE: A multiple sequence alignment method with reduced time and space complexity. BMC Bioinformatics 5:113. https://doi.org/10.1186/1471-2105-5-113

62. Larsson A (2014) AliView: a fast and lightweight alignment viewer and editor for large datasets. Bioinformatics 30:3276–3278

63. Kalyaanamoorthy S, Minh BQ, Wong TKF, et al (2017) ModelFinder: Fast model selection for accurate phylogenetic estimates. Nature Methods 14:587–589. https://doi.org/10.1038/nmeth.4285

64. Penev PI, Alvarez-Carreño C, Smith E, et al (2021) TwinCons: Conservation score for uncovering deep sequence similarity and divergence. PLoS Computational Biology 17:e1009541. https://doi.org/10.1371/journal.pcbi.1009541

65. Wheeler TJ, Clements J, Finn RD (2014) Skylign: A tool for creating informative, interactive logos representing sequence alignments and profile hidden Markov models. BMC Bioinformatics 15:7. https://doi.org/10.1186/1471-2105-15-7

66. Lampropoulos A, Sutikovic Z, Wenzl C, et al (2013) GreenGate---a novel, versatile, and efficient cloning system for plant transgenesis. PLoS One 8:e83043. https://doi.org/10.1371/journal.pone.0083043

67. Perlaza K, Toutkoushian H, Boone M, et al (2019) The mars1 kinase confers photoprotection through signaling in the chloroplast unfolded protein response. eLife 8:1–36. https://doi.org/10.7554/eLife.49577

68. Bechtold N, Pelletier G (1998) In planta Agrobacterium-mediated transformation of adult Arabidopsis thaliana plants by vacuum infiltration. Methods in molecular biology (Clifton, NJ) 82:259–266. https://doi.org/10.1385/0-89603-391-0:259

69. Schindelin J, Arganda-Carreras I, Frise E, et al (2012) Fiji: an open-source platform for biological-image analysis. Nature methods 9:676–682. https://doi.org/10.1038/nmeth.2019

70. Marley J, Lu M, Bracken C (2001) A method for efficient isotopic labeling of recombinant proteins. Journal of biomolecular NMR 20:71–75. https://doi.org/10.1023/a:1011254402785

71. Schmidt U, Weigert M, Broaddus C, Myers G (2018) Cell detection with star-convex polygons. Lecture Notes in Computer Science (including subseries Lecture Notes in Artificial Intelligence and Lecture Notes in Bioinformatics) 11071 LNCS:265–273. https://doi.org/10.1007/978-3-030-00934-2_30

72. Delaglio F, Grzesiek S, Vuister GW, et al (1995) NMRPipe: a multidimensional spectral processing system based on UNIX pipes. Journal of biomolecular NMR 6:277–293. https://doi.org/10.1007/BF00197809

73. Skinner SP, Fogh RH, Boucher W, et al (2016) CcpNmr AnalysisAssign: a flexible platform for integrated NMR analysis. Journal of biomolecular NMR 66:111–124. https://doi.org/10.1007/s10858-016-0060-y

74. R Core Team (2021) R: A Language and Environment for Statistical Computing

