## Supplementary material for "Shuffled ATG8 interacting motifs form an ancestral bridge between UFMylation and C53-mediated autophagy": Analyze_Agarose_beads_v2.ijm

```
run("Close All");
inDir = getDirectory("Directory Containing Files "); 
ModelPath = inDir + "Lorenzo_Widefield_new2.zip";
run("Options...", "iterations=1 count=1 black do=Nothing");
//print(ModelPath);
ModelPathNew=replace(ModelPath,"\\","/");
//print(ModelPathNew);
//Exit()
//E:\\\\Penninger\\\\TestforMacro\\\\Vanja_RGB_2x_downsample.zip
//outDir = getDirectory("Folder for Export");
fileList = getFileList(inDir);
print(fileList.length);
run("Clear Results");

NucleiOutsideLarge = newArray(fileList.length);

Count = 0;

///Set the minimum and maximum area threshold
AreaThreshMin = 100; 
AreaThreshMax = 100000;

//Set Channel to perform the deep learning
MeasureChannel = 2;

//Set Channel for counting cells
CountCH = 2;

//Set downsampling factor, 1 is original size, 2 is half size etc.
//Downsample = 1;


//Define detection of Cells on filtered or unfiltered image. "0" means no filter.

BandpassFilter = 0;


count=1;

setBatchMode(true);

for (n=0; n<fileList.length-1; n++){
	////Cut Imagename
	inFileCut = lengthOf(fileList[n])-4;
	//print(inFileCut);
	saveName=substring(fileList[n],0,inFileCut);
	Name=fileList[n];
	file = inDir + fileList[n];
	//print(file); 
	//print(Name);
	//exit

	////Find mrxs files
	Run = 0;
	matching=(endsWith(Name, ".czi"));
	if (matching==1) Run=1;
	else {};
	matching=(endsWith(Name, ".csv"));
	if (matching==1) Run=0;
	else {};	
	matching=(endsWith(Name, ".xls"));
	if (matching==1) Run=0;
	else {};	
	matching=(endsWith(Name, "segment.tif"));
	if (matching==1) Run=0;
	else {};	

	print(Run);
	if (Run == 1){
	
		setBatchMode(false);
		run("Close All");
		run("Clear Results");
		roiManager("reset");
		
		run("Bio-Formats Importer", "open="+file+" color_mode=Default rois_import=[ROI manager] view=Hyperstack stack_order=XYCZT");
		run("Duplicate...", "title=CH"+MeasureChannel+" duplicate channels="+MeasureChannel+"");
	
		//././././ Perform deep Learning

		run("Command From Macro", "command=[de.csbdresden.stardist.StarDist2D], args=['input':'CH"+MeasureChannel+"', 'modelChoice':'Model (.zip) from File', 'normalizeInput':'true', 'percentileBottom':'1.0', 'percentileTop':'99.8', 'probThresh':'0.25', 'nmsThresh':'0.4', 'outputType':'Both', 'modelFile':'"+ModelPathNew+"', 'nTiles':'1', 'excludeBoundary':'2', 'roiPosition':'Automatic', 'verbose':'false', 'showCsbdeepProgress':'false', 'showProbAndDist':'false'], process=[false]");
		//run("Command From Macro", "command=[de.csbdresden.stardist.StarDist2D], args=['input':'slide-2020-10-13T14-21-25-R1-S2-Annotation10_stack.tif_CenterSlice.tif (RGB)', 'modelChoice':'Model (.zip) from File', 'normalizeInput':'true', 'percentileBottom':'1.0', 'percentileTop':'99.8', 'probThresh':'0.5', 'nmsThresh':'0.25', 'outputType':'Both', 'modelFile':'"+ModelPathNew+"', 'nTiles':'1', 'excludeBoundary':'2', 'roiPosition':'Automatic', 'verbose':'false', 'showCsbdeepProgress':'false', 'showProbAndDist':'false'], process=[false]");
		setBatchMode(true);
		//run("Set Measurements...", "mean standard modal min median redirect=None decimal=3");
		run("Set Measurements...", "area mean standard modal min shape median redirect=None decimal=3");
		run("Line Width...", "line=5");
		Number = roiManager("count");

		run("Clear Results");
		run("Duplicate...", "title=Mask duplicate channels=1");
		run("8-bit");
	
		run("Grays");
		run("Select All");
		run("Clear", "slice");

		for(i=0; i<Number; i++){
			//selectWindow(Name);
			roiManager("select", i);
			run("Enlarge...", "enlarge=2 pixel");
			//roiManager("Update");
			run("Invert");
		}
		run("Watershed");
		run("Kill Borders");
		close("Mask");
		
		////Remove border objects and appy size thresholds
		
		selectWindow("Mask-killBorders");
		ThreshROI("Mean", 200, ">");
		
		ThreshROI("Round", 0.96, ">");
		selectWindow("Mask-killBorders");
		ThreshROI("Area", AreaThreshMin, ">");
		ThreshROI("Area", AreaThreshMax, "<");

		//nRoi = roiManager("count");

		rename("Mask");
		run("32-bit");
		setAutoThreshold("Default dark");
		//run("Threshold...");
		run("NaN Background");
		run("Divide...", "value=255.000");
		selectWindow(Name);
		run("Duplicate...", "title=CH1 duplicate channels=1");
		imageCalculator("Multiply create 32-bit", "CH1","Mask");
		rename("Measure");
		run("Clear Results");
		
		Number = roiManager("count");		
		print("Number = "+Number);
		if (Number == 0){
			selectWindow("CH2");
			run("Enhance Contrast", "saturated=0.35");
			wait(50);
			saveAs("jpg", inDir + Name + "_BF");
			//Exit()
			selectWindow("Measure");
			saveAs("jpg", inDir + Name + "_Fluorescence");	
		}

		else{
			for(i=0; i<Number; i++){
				selectWindow("Measure");
				roiManager("select", i);
				run("Area to Line");
				run("Line to Area");
				roiManager("Rename", i+1);
				//Stack.setChannel(1);
				//run("Measure");
				roiManager("Update");
			}
			roiManager("deselect");
			roiManager("measure");

			saveAs("Results", inDir + Name + "_Results.xls");
			selectWindow("CH2");
			run("Enhance Contrast", "saturated=0.35");
			wait(50);
			run("From ROI Manager");
			run("Flatten");
			saveAs("jpg", inDir + Name + "_BF");
			//Exit()
			selectWindow("Measure");
			run("From ROI Manager");
			run("Flatten");
			saveAs("jpg", inDir + Name + "_Fluorescence");
		}
		//Exit()
	}
	//print(count);
	count++;
}

//run("Analyze Particles...", "display clear add");
run("Clear Results");
//roiManager("deselect");
//roiManager("measure");

ThreshROI("Area", 30, ">");
ThreshROI("StdDev", 5000, ">");

function ThreshROI(Parameter, Value, Sign){
	run("Clear Results");
	run("Select None");
	roiManager("deselect");
	roiManager("measure");

	//sortArray = newArray(nResults);
	res=0;
	numb=0;
	nRoi = roiManager("count");
	if(Sign == ">"){
		for(i=0; i<nResults; i++){
			print("Number = "+numb);
			y = getResult(Parameter, i);
			print(y);
			if (y < Value){
				roiManager("select", numb);
				roiManager("delete");
				print("ROI = "+numb);
				//exit
				numb--;
				
			}
			else{};			
			numb++;
			//nRoi = roiManager("count");
			
		}
	}
		if(Sign == "<"){
		for(i=0; i<nResults; i++){
			print("Number = "+numb);
			y = getResult(Parameter, i);
			print(y);
			if (y > Value){
				roiManager("select", numb);
				roiManager("delete");
				print("ROI = "+numb);
				//exit
				numb--;
				
			}
			else{};			
			numb++;
			//nRoi = roiManager("count");
			
		}
	}
	
	roiManager("Show All");
	run("Select None");
}


run("Close All");
Exit()


function Exit(){
	setBatchMode("show all");
	exit();
}
```
